# Predicting neurodevelopmental outcomes in children with perinatal HIV using a novel machine learning algorithm

**DOI:** 10.1101/632273

**Authors:** Robert Paul, Kyu Cho, Claude Mellins, Kathleen Malee, Reuben Robbins, Stephen Kerr, Jiratchaya Sophonphan, Neda Jahanshad, Linda Aurpibul, Kulvadee Thongpibul, Pope Kosalaraksa, Suparat Kanjanavanit, Chaiwat Ngampiyaskul, Wicharn Luesomboon, Jurai Wongsawat, Saphonn Vonthanak, Kea Chettra, Tulathip Suwanlerk, Victor Valcour, Lila Balla, Paola M. Garcia-Egan, Rebecca Preston, Jintanat Ananworanich, Thanyawee Puthanakit, on behalf of the PREDICT Study Group

## Abstract

**Background:** A subset of children with perinatal HIV (pHIV) experience long-term neurocognitive symptoms despite treatment with antiretroviral therapy. However, predictors of neurocognitive outcomes remain elusive, particularly for children with pHIV residing in low-to-middle income countries. The present study utilized a novel data analytic approach to identify clinically-relevant predictors of neurocognitive development in children with pHIV.

**Methods:** Analyses were conducted on a large repository of longitudinal data from 285 children with pHIV in Thailand (n=170) and Cambodia (n=115). Participants were designated as neurocognitively resilient (i.e., positive slope; n=143) or at risk (i.e., negative slope; n=142) according to annual performances on the Beery-Buktenica Developmental Test of Visual-Motor Integration over an average of 5.4 years. Gradient-boosted multivariate regression (GBM) with 5-fold cross validation was utilized to identify the optimal combination of demographic, HIV disease, blood markers, and emotional health indices that predicted classification into the two neurocognitive subgroups. Model performance was assessed using Receiver Operator Curves and sensitivity/specificity.

**Results:** The analytic approach distinguished neurocognitive subgroups with high accuracy (93%; sensitivity and specificity each > 90%). Dynamic change indices and interactions between mental health and biological indices emerged as key predictors.

**Conclusion:** Machine learning-based regression defined a unique explanatory model of neurocognitive outcomes among children with pHIV. The predictive algorithm included a combination of HIV, physical health, and mental health indices extracted from readily available clinical measures. Studies are needed to explore the clinical relevance of the data-driven explanatory model, including potential to inform targeted interventions aimed at modifiable risk factors.

## Introduction

A subset of children with perinatal-acquired human immunodeficiency virus (pHIV) exhibit neurocognitive difficulties [1-2] despite use of antiretroviral therapy (ART) [3]. Initial findings from a prospective investigation of newborns with pHIV [4] described better neurocognitive outcomes at age 1 when ART was started within the first 30 days of life. However, recent analyses indicate that the neurocognitive advantages attributed to initiation of ART during the immediate postnatal period are not sustained for many individuals in later childhood [5]. Similar findings have been reported in separate cohorts of older children with pHIV who initiated suppressive ART within weeks of birth [6-7], as well as subgroups of children who initiated ART after surviving the first year of life without significant immune compromise [8-13].

Predictive models of neurocognitive outcomes in children with pHIV remain elusive, especially for individuals residing in resource-restricted environments where most of the global population of children with pHIV reside. Traditional data analytic methods are not well suited to discover mechanisms that underlie complex clinical phenotypes such as neurocognitive development [see 14 for review]. Common statistical models are inherently restricted by statistical assumptions (e.g., normality), redundancy across variables (e.g., multicollinearity), and reliance on conservative significance thresholds to reduce errors resulting from multiple comparisons [14]. Additionally, traditional methods, such as logistic regression, require a priori selection of predictor variables despite limited insights into the underlying data structure [14]. This approach is appropriate for confirmation testing, but highly punitive for work aimed at scientific discovery.

Advanced data science methods such as machine learning offer a promising alternative to detect latent variables that underlie complex clinical phenotypes such as HIV [14-21]. In studies of structural neuroimaging, Wade et al. [19] achieved 72% accuracy classifying adults as either HIV-infected or HIV-uninfected, and Zhang et al. [20] achieved 85% accuracy discriminating older adults with HIV from age-matched adults with early Alzheimer’s disease. Most recently, Ogishi et al. [21] trained a classifier using viral signatures to identify adults with clinically-defined severe neurocognitive impairment (HAD) versus individuals with no neurocognitive impairment. In this study, supervised machine learning achieved 90% predictive accuracy.

The studies above utilize cross-sectional data to develop a data-driven algorithm for group classification. Only one study [21] examined utilized advanced methods to explore the underlying mechanisms, but the study solely focused on viral signatures. No studies have integrated demographic, HIV disease variables, routine laboratory indices, and psychosocial variables to establish a robust predictive algorithm for neurocognitive sequelae of HIV. Further, no studies have leveraged the methodological advantages of machine learning to establish a data-driven classifier of neurocognitive outcomes in children with pHIV. This represents an important gap in the literature considering that pediatric populations are likely to benefit from evidence-based methods to enrich neurocognitive outcomes.

The current study provides a foundation to address this gap in the literature. We used unsupervised machine learning to explore intrinsic patterns of bio-behavioral data within a large, high dimensional longitudinal dataset of 285 children with pHIV from Thailand and Cambodia [8-9]. Here we leveraged the computational strengths of supervised machine learning to establish a data-driven algorithm of neurocognitive risk versus resilience among children with pHIV, and to discover novel predictors of neurocognitive risk stratification using common clinical measures.

## Methods

### Study Design

Archival data were analyzed from 285 Thai (n=170) and Cambodian (n=115) youth with pHIV enrolled in the Pediatric Randomized to Early vs. Deferred Initiation in Cambodia and Thailand (PREDICT) clinical trial (clinicaltrials.gov identification: U19AI53741) [8-9]. Enrollment for PREDICT began in 2006, before global treatment guidelines recommended ART at the time of diagnosis. The PREDICT study enrolled children with pHIV with CD4 % between 15-25% and no history of ART. Participants were then randomized to start ART immediately or when CD4 declined to 15%. Mortality and the frequency of AIDS-defining illnesses were evaluated 144 weeks from the date of enrollment. A neuro substudy began one year after the start of the main PREDICT trial (MH#). Published outcomes report no differences in survival, AIDS-defining illnesses, or neurocognitive outcomes between the two treatment arms [8-13]. Most participants enrolled in PREDICT subsequently enrolled in an observational follow-up investigation (RESILIENCE; MH# R01MH102151).

The present study included children ages 2-14 years at the time of the first neurocognitive assessment (consistent with the minimum age requirement for the primary outcome measure described below). As defined by the parent PREDICT protocol, individuals were excluded if they reported a history of brain infection, neurological or psychiatric disorder, congenital abnormalities; previous use of immunomodulators within 4 weeks of enrollment; baseline absolute neutrophil count < 750 cells/μL, hemoglobin < 7.5 g/dL; baseline platelet counts < 50,000 per μL, or alanine aminotransferase > 4 times the upper limit of normative values. Approval was obtained by participating Institutional Review Boards. Caregivers provided informed consent and assent was obtained from children older than 7 years of age. Demographic and clinical characteristics are provided in **Table 1**.

**Table 1.**
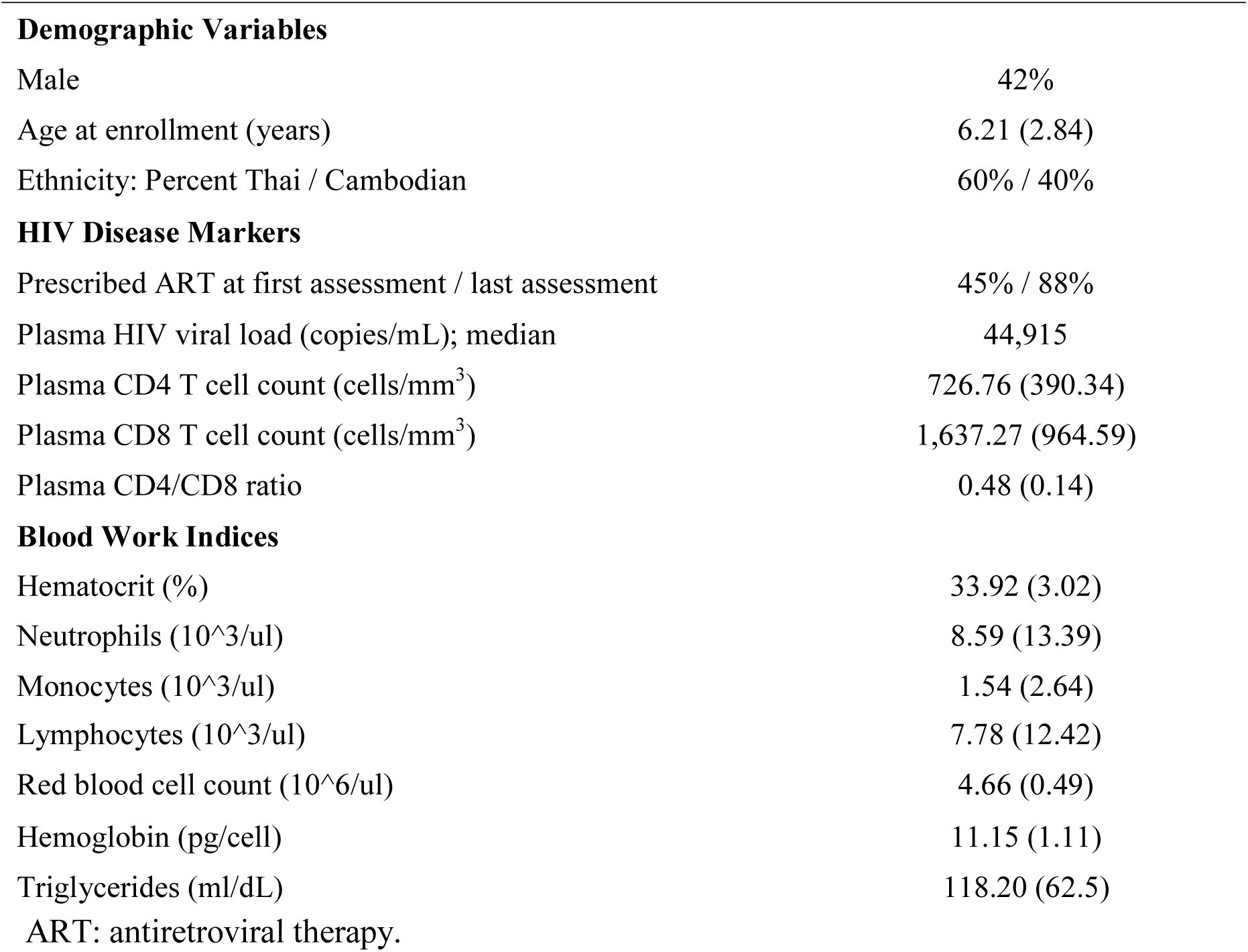
Sample demographics and clinical variables at enrollment.

#### Assessment of neurocognitive development

The Beery-Buktenica Developmental Test of Visual-Motor Integration, Fourth Edition (BVMI) [22] served as the measure of neurocognitive development. The BVMI is a standardized test of graphomotor construction of 30 increasingly complex line drawings that are presented in sequential order. The test has been utilized extensively in international studies of pediatric neurodevelopment, including Thai and Cambodian children with pHIV [8-13]. The BVMI was developed for individuals from ages 2-90 years from diverse environmental, educational, and linguistic backgrounds [22]. Performance is strongly related to chronological age [22-23]; longitudinal studies of normative samples reveal an average increase in the total score of about 15 points from age 2 to age 20 years [23]. The BVMI was administered at enrollment (baseline), twice per year during the PREDICT trial, and then annually thereafter. The average duration of follow-up for the current analysis was 5.4 years.

Neurocognitive trajectories on the BVMI were quantified as the average percent change in raw score performance from baseline to the final test visit (i.e., positive vs. negative slopes) using all available performance data. Participants were designated as neurocognitively resilient (blue trajectories: n=145) or at-risk (red trajectories: n=140) based on a within sample comparison (upper/lower halves of the sample distribution; **Figure 1**). This binary designation optimized the number of observations to train the machine learning classifier while minimizing potential bias associated with various definitions of clinically-relevant neurocognitive impairment.

**Figure 1.**
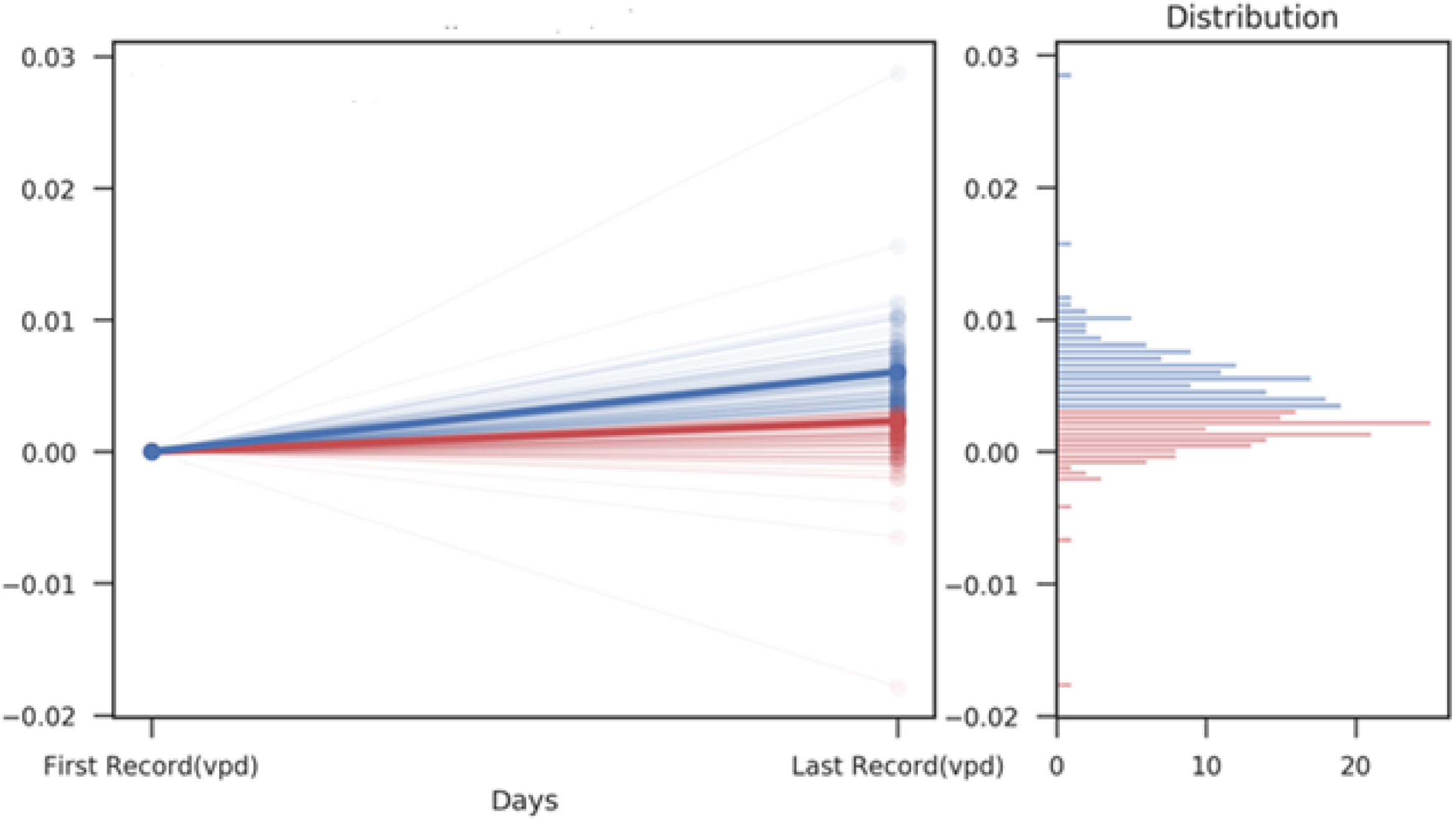
Trajectories (slopes) representing the upper and lower halves of the performance continuum on the BVMI over an average of 5.4 years. Neurocognitively resilient group (blue) and neurocognitively at risk group (red).

#### Candidate predictors of neurocognitive development

The major categories of candidate predictors (i.e., input features) included: 1) *demographic variables* (age, sex, family income, etc.); 2) *HIV disease indices* (blood-derived CD4 T cell count, CD8 T cell count, viral load, etc.); 3) *routine blood markers* (complete blood count, glucose levels, triglycerides, etc.); and 4) *emotional health indices* (component and domain scales from the Child Behavior Checklist-Caregiver version (CBCL) [24]. Caregivers of participants completed age-appropriate versions of the CBCL (age 2 to 5 years or age 6 years and over), translated into Thai and Khmer. Raw scores were converted to standardized scores (T scores) to allow comparison to published studies in this population [8-13].

#### Machine learning approach

Our predictive algorithm was built using gradient boosted multivariate regression (GBM), a form of machine learning. GBM is based on random forest decision tree analysis, which utilizes recursive partitioning to iteratively populate homogeneous predictors [25]. The final output from the decision tree is represented mathematically as the lowest possible sum of squares. We set the default number of decision trees at 10,000 and shrinkage at 0.005. Prediction accuracy was calculated by transforming the raw output into a probability score using the sigmoid function (1/(1+e^(-x)). The assigned class label employed a 0.5 decision boundary for evaluation. Gradient descent was applied to minimize prediction errors. Performance indicators included the area under the curve (AUC), sensitivity, specificity, and the F1 score (i.e., harmonic balance between sensitivity and specificity). Calculations for these indices are described in detail elsewhere [26]. AUC interpretation followed standard convention (poor =<0.6, fair =0.7, good=0.8; excellent >0.8) [26]. Stability of the final model was examined using 5-fold cross-validation.

#### Machine learning feature selection

The machine learning analysis was conducted in Python. Feature selection was conducted using an in-house program based on SciKit [27] and PDPBox [28]. Synthetic data elements (e.g., participant identification number) were not included in the machine learning analysis but all biologically plausible variables were examined without a priori bias related to predictive relevance. For each predictive variable, we considered multiple permutations of central tendency (e.g., average, max value), dispersion (e.g., standard deviation, percent variability) and change over time (e.g., average value per day, minimum percent change per day). Polynomial interactions were explored up to 3-way interactions. We selected 3-levels based on prior studies demonstrating 3-way interactions between environmental factors, biometrics, and psychosocial indices [29]. For behavioral inputs, feature clusters included both composite and subscale components (e.g., CBCL Total Score and CBCL Affective scale score) if the combination improved classification accuracy (i.e., the variables were not duplicative inputs).

#### Post-hoc analysis of data-driven predictors

The top 25 predictors of neurocognitive risk vs. resilience were compared using R. Mean values from the first and last visits were contrasted using Kruskal-Wallis test (p < 0.05), adjusted for multiple comparisons. Additionally, we conducted a standard logistic regression in R to establish a benchmark using traditional statistical analyses. In accord with standard convention, the maximum number of independent variables in the logistic regression was governed by the sample size, allowing for a total of 16 variables (four independent variables and each of the two-way interactions; all from baseline). The four independent variables included average family income, CD4/CD8 T-cell ratio, HIV viral load, and CBCL Total Score. These variables represented demographic, immunological, disease severity, and emotional health indices, respectively. The logistic regression was modeled using mean values versus indices informed by the machine learning results. This approach provided an opportunity to compare the results of machine learning to a traditional logistic regression. Fisher’s Exact test [30] examined the classification accuracy obtained by logistic regression and the GBM.

## Results

### Demographic and clinical variables

At baseline, 157 (55%) participants were prescribed ART. First-line treatment included zidovudine, lamivudine, and nevirapine. One individual was diagnosed with esophageal candidiasis within the first month of enrollment; no other AIDS-defining illnesses were observed in either treatment arm. Between the time period of enrollment and the last neurocognitive assessment, 124 (43%) participants randomized to the deferred treatment arm exhibited a decline in CD4% that met threshold for ART initiation (i.e., < 15%). These individuals immediately began ART. The remaining 33 (11%) children did not experience a decline in CD4 T cell count < 15% before the final BVMI assessment; these participants remained untreated throughout the follow-up period included in the current analysis.

### Machine learning classification results

The GBM classifier of neurocognitive trajectories achieved high accuracy (93%), with 92% sensitivity and 94% specificity (F1 score= 0.93). The average AUC was 95% (range 93-99%). The final algorithm included 25 predictive features comprised of 54 individual data elements. Receiver Operator Curve (AUC) results with 5-fold cross validation are depicted in **Figure 2**.

**Figure 2.**
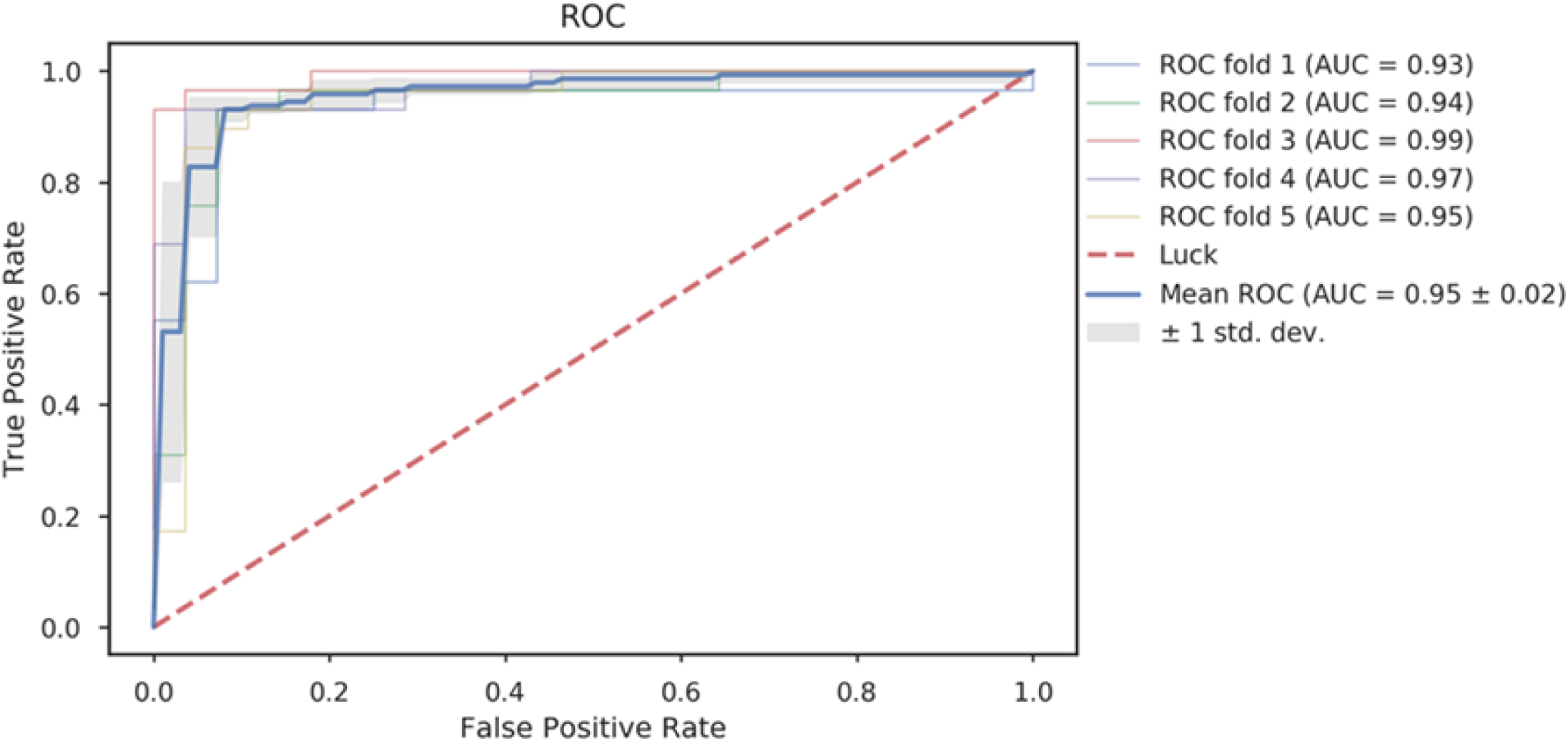
Area Under the Curve achieved by the machine learning model with 5-fold cross validation. AUC range .93-99; average = .95.

The 25 classification features that emerged from the machine learning analysis are presented in **Figure 3** in rank order of classification relevance (see Supplement for box plots). The full algorithm included a collection of polynomial interactions (combinatorial features) and individual classifiers. Values of change, dispersion, and central tendency were represented in the algorithm. The top 10 classification features are described below. The mathematical operations underlying the polynomial interactions are provided in the Supplement.

**Figure 3.**
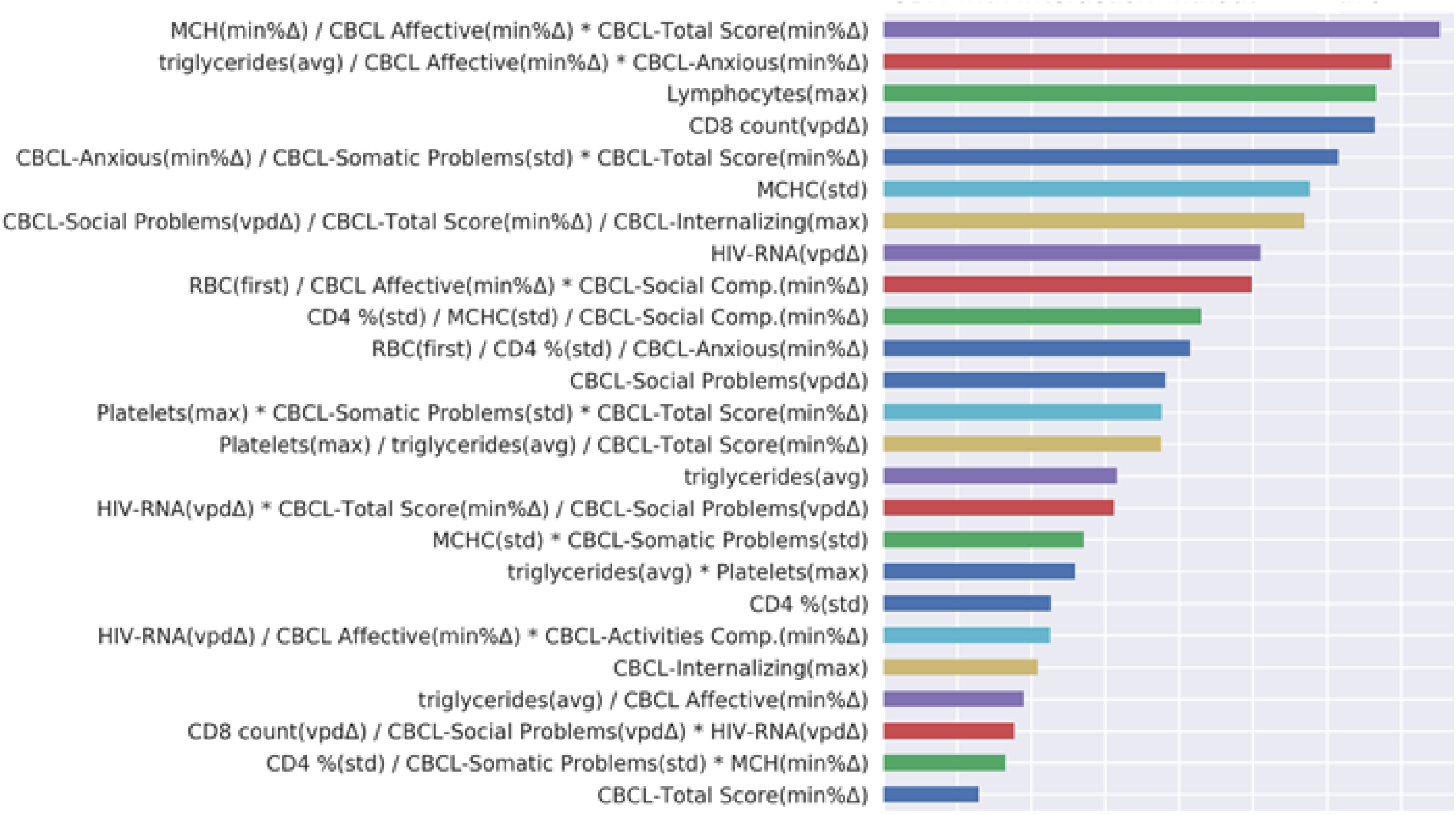
Feature importance (by rank order). Abbreviations: avg, average value; first, first recorded value; max, maximum recorded value; min % Δ, minimum percent change; vpd Δ, value per day change. Triglycerides (md/dL); HIV-RNA (copies/mL); MCHC, Mean corpuscular hemoglobin concentration (g/dL); Platelets (10^3/ul); RBC, Red blood cell (10^6/ul); Lymphocytes (10^3/ul); *Child Behavior Checklist Values (CBCL*): Total score; Affective Problems; Somatic Complaints; Social Problems; Anxious Scale; Internalizing Scale, Social Competency; Activities Competence. All CBCL values represent T scores.

#### Combinatorial Features (1 through 6)

**Feature 1)** higher hemoglobin, lower CBCL Affective scale, and lower CBCL Total Score (each as percent change); **Feature 2)** lower average triglyceride level, and lower CBCL Affective and CBCL Anxious scales (both as minimum percent change); **Feature 3)** lower CBCL Anxious scale (minimum percent change), lower CBCL Somatic Problems scale standard deviation, and lower CBCL Total Score (minimum percent change); **Feature 4)** lower CBCL Social Problems scale (value per day), lower CBCL Total Score (minimum percent change), and lower max change in CBCL Internalizing scale; **Feature 5)** higher total erythrocyte count, lower CBCL Affective scale (minimum percent change), and lower CBCL Social Competence scale (minimum percent change); and **Feature 6)** lower percent variability in CD4 cell count, lower erythrocyte count standard deviation, and lower CBCL Social Competence scale (minimum percent change).

#### Discrete Features (A through D)

**Feature A)** higher max lymphocyte count; **Feature B)** higher plasma CD8 T count (value per day); **Feature C)** lower hemoglobin standard deviation; and **Feature D)** lower plasma HIV viral load (value per day). See Table 2 for contrasts between groups on the top 10 features.

**Table 2.**
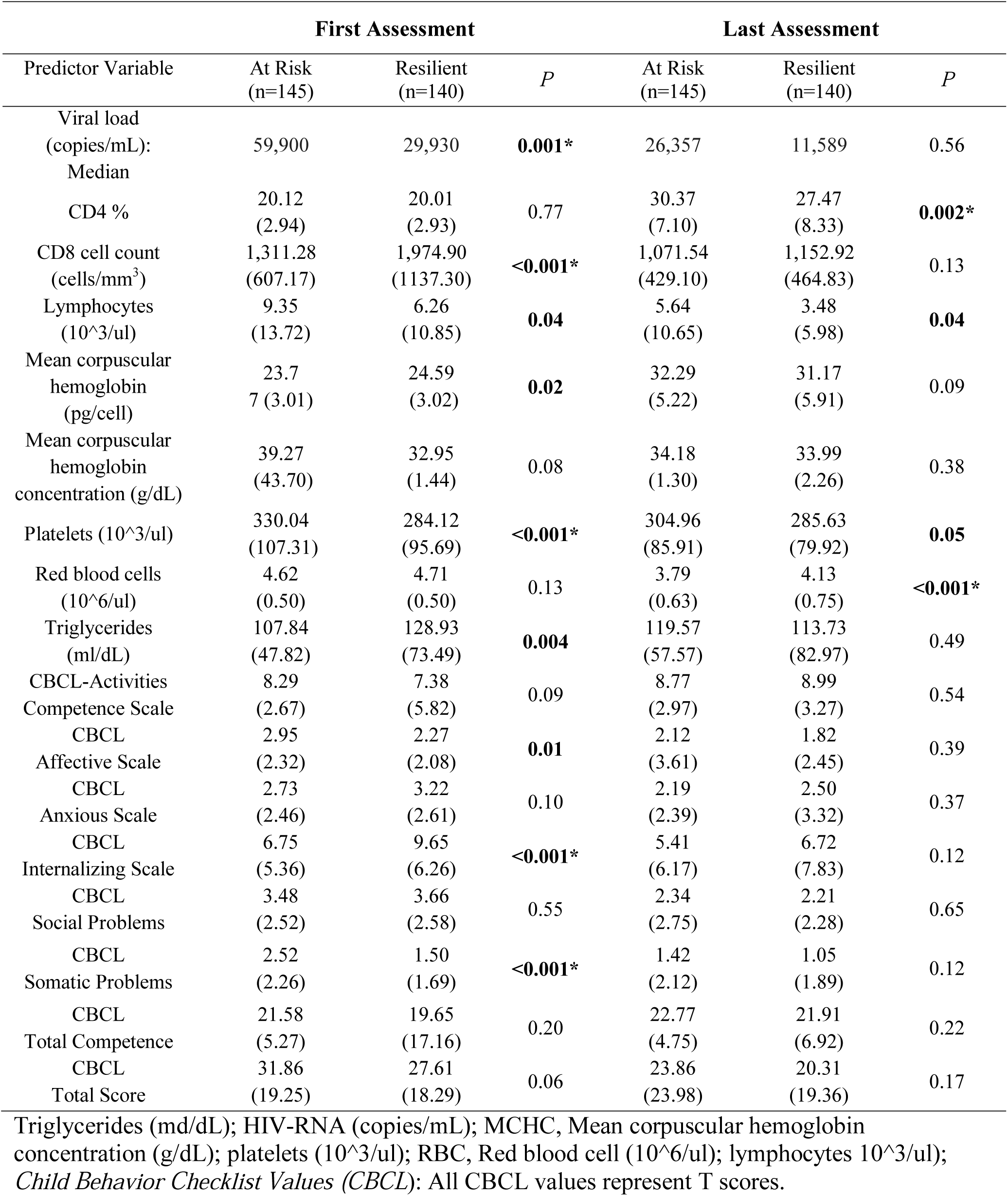
Comparisons between the two neurocognitive risk groups on variables selected by the machine learning analysis.

### Posthoc logistic regression

The traditional logistic regression analysis using a subset of candidate predictors was nonsignificant (45% (X^2^(10) = 44.6, log likelihood: 175.20; pseudo R^2^: 0.11; Table S2). Total classification accuracy was 45% (compared to 93% using the machine learning approach). Variables retained in the final logistic regression model included two interaction terms: 1) HIV viral load and CBCL Total Score; and 2) HIV viral load and family income. The difference in classification accuracy between the GBM algorithm (93%) and the logistic regression model (45%) was statistically significant (p =0.035; Fisher’s Exact test).

## Conclusions

Using supervised machine learning and intelligent feature creation and selection on highly dimensional data repository, we established a robust classifier of divergent neurocognitive trajectories in children with pHIV. The classifier yielded an average accuracy of group membership that was more than two times better than results obtained using standard logistic regression. Importantly, the machine learning method identified novel risk factors for suboptimal neurocognitive outcomes in this cohort of youth with pHIV, including numerous multi-level interactions between mental health and biological indices. The final classification algorithm was comprised of 25 predictive features derived from 54 data elements. The study is the first to integrate a large number of relevant risk factors to establish a predictive algorithm for neurocognitive outcomes in children with pHIV residing in resource-restricted environments. Study findings demonstrate the potential for data-driven methods to generate new insights into the underlying mechanisms associated with divergent neurocognitive outcomes in children with pHIV.

Prior investigations aimed at unraveling the neuropathogenic model of pHIV have often leaned on the adult literature to select candidate mechanisms. This approach assumes that neurocognitive symptoms in pediatric and adult populations result from the same underlying processes. However, the scientific basis for this assumption is weak. Children with pHIV are typically infected in utero, when the brain is undergoing rapid, dynamic changes in neuronal architecture and functional organization [31-34]. After birth, environmental pressures, including living with a chronic medical disease, moderate brain development and expression of brain-behavior phenotypes [35-38]. Many children infected with HIV also face chronic stress associated with poverty (food security and malnutrition, limited education, family and social support disruption; [6-13]). Stigma related to HIV and psychological challenges created by HIV disclosure among sexually active adolescents and young adults contribute to the burden of early life adversities in this vulnerable population [39].

Previous machine learning studies of brain integrity in people living with HIV have exclusively targeted adult samples [19-21]. Our study is the first to establish a machine learning classifier of neurocognitive performance in children with pHIV. Our GBM approach differentiated individuals designated as neurocognitively resilient vs. at risk. Unlike studies in adult samples [21], we focused on children without evidence of severe neurocognitive impairment and without a history of brain trauma. Differentiation of individuals with less severe neurocognitive impairment from normative performance requires analytic methods capable of integrating more input variables than what can be achieved with traditional statistics. Results from this study underscore this point, as the machine learning algorithm performed more than two-fold better than logistic regression in classifying individuals with divergent neurocognitive outcomes. Our decision to include individuals without overt neurocognitive impairment is consistent with work in other populations in which predictive modeling targets the prodromal stages of neurocognitive dysfunction [40-41]. Early detection of neurocognitive risk is especially important for pediatric populations who can benefit from informed educational/therapeutic services.

Our set of input features included a broad range of typical (e.g., means, max), and less common (minimum percent change) metrics without a priori bias as to which variables were most relevant to the outcomes. Interestingly, only one predictive feature (baseline triglyceride level) was defined by a typical average value; all other predictors represented change (i.e., temporal patterns) or dispersion metrics. The poor sensitivity of average values was borne out by the logistic regression model and by the between-group contrasts from the first and last visits, neither of which identified a consistent profile of risk. By contrast, temporal patterns discovered by the machine learning analysis were salient predictive features, representing 75% of the final algorithm (**Supplement Figure 2**). These findings reinforce results from prior studies demonstrating poor stability and generalizability of predictive models built from cross-sectional data [42-44]. Prospective research is resource-intense and computationally challenging, but longitudinal cohort studies such as PREDICT are necessary to develop accurate longitudinal models (e.g., time series forecasting) of neurodevelopment.

Another important result from this study is the synergy observed between mental and biological markers of health and HIV (e.g., triglyceride levels, erythrocyte count, viral load, CD4 count). Mental health emerged as an important element in 28% of the algorithm (**Supplemental Table 1**). Consistent/stable scores reported on the CBCL Anxiousness, Affective, Somatic Problems, and Social Competence subscales interacted with higher erythrocyte count, lower triglyceride level, CD4, greater degree of HIV RNA reduction, and higher platelet counts. Whether the expression of mental health symptoms precedes, follow, or emerges concurrently with change in these biometrics cannot be ascertained from the current study. However, the observation that > 50% of the top 10 classifiers were comprised of multi-level interactions involving mental health emphasize the potential clinical value of synergies across predictors rather than cumulative models of risk stratification.

Interestingly, several polynomial interactions were comprised of composite scores and inter-related subscales (e.g., CBCL Total Score and CBCL Affective scale). While these variables are inherently intertwined, each contribute a degree of explanatory variance in neurocognitive trajectories. Machine learning methods are well suited to examine these interdependencies, albeit this often occurs at the expense of clear relevance to clinical meaning. Prior work describes the “black box dilemma” [14] of machine learning algorithms that favor classification accuracy over clinical interpretation. Deep learning algorithms, similar to the approach utilized in this study, offer preliminary insights into the underlying mechanisms, but follow-up work with item-level analyses are needed.

Limitations of the study merit discussion. Overfitting is a concern with any predictive model, including data-driven methods such as machine learning. The specific approach utilized in this study is relatively robust to overfitting and other sources of error such as unbalanced group designs, and low base rates of the targeted outcome [14]. Nevertheless, we would expect some loss in predictive accuracy when the algorithm is applied in an independent cohort, particularly since participants enrolled in PREDICT represent a unique group of individuals with pHIV who survived the first year of life without ART. As such, it is unlikely that the algorithm generated for this cohort would apply equally to other populations of pHIV. Finally, as a preliminary study, we explored numerous polynomial interactions and various metrics beyond central measures of tendency. Additional studies are needed to examine the clinical relevance of predictors defined by nontraditional metrics.

In summary, the present study provides a critical first step towards establishing a data-driven predictive model of neurocognitive outcomes in children with pHIV. We cannot fully interpret the clinical relevance of the novel mechanisms identified by the machine learning analysis, but the results provide a compelling argument that mental health interacts with numerous risk factors to govern neurodevelopmental outcomes in children with pHIV. Ideally, these results will guide new data-driven investigations and inform the development of interventions aimed at modifiable risk factors/facilitators of overall health and wellness in children with pHIV.

## Authors’ contributions

Study conceptualization and design: (RP, KC, CM, KM, RR, SK, LS, VV, JA, TP) Data analysis: (KC, JS, LB, PG); Substantial data collection: (JS, LA, KT, PK, CN, WL, JW, SV, KC, TS); Substantial writing contribution: (RP, KC, CM, KM, RR, SK, LS, VV, JA, TP)

## Acknowledgement

We are grateful to the children and caregivers for their participation in this study. The PREDICT study was sponsored by the National Institute of Allergy and Infectious Disease (NIAID), Grant number U19 AI053741, Clinical trial.gov identification number NCT00234091. Antiretroviral drugs for PREDICT were provided by ViiV Healthcare (AZT, 3TC), Boehringer Ingelheim (NVP), Merck (EFV), Abbott (RTV) and Roche (NFV). The neurodevelopment work was supported by initial funds by the Eunice Kennedy Shriver National Institute of Child Health and Human Development and the National Institute of Mental Health, and later funded by R01MH089722 (Valcour) and R01MH) 102151Ananworanich(and Puthanak.

The PREDICT Study Group

HIV Netherlands Australia Thailand (HIV-NAT) Research Collaboration, Thai Red Cross AIDS Research Center, Bangkok, Thailand; Kiat Ruxrungtham, Jintanat Ananworanich, Thanyawee Puthanakit, Chitsanu Pancharoen, Torsak Bunupuradah, Stephen Kerr, Theshinee Chuenyam, Sasiwimol Ubolyam, Apicha Mahanontharit, Tulathip Suwanlerk, Jintana Intasan, Thidarat Jupimai, Primwichaya Intakan, Tawan Hirunyanulux, Praneet Pinklow, Kanchana Pruksakaew, Oratai Butterworth, Nitiya Chomchey, Chulalak Sriheara, Anuntaya Uanithirat, Sunate Posyauattanakul, Thipsiri Prungsin, Pitch Boonrak, Waraporn Sakornjun, Tanakorn Apornpong, Jiratchaya Sophonphan, Ormrudee Ritim, Nuchapong Noumtong, Noppong Hirunwadee, Chowalit Phadungphon, Wanchai Thongsee, Orathai Chaiya, Augchara Suwannawat, Threepol Sattong, Niti Wongthai,Kesdao Nantapisan, Umpaporn Methanggool, Narumon Suebsri, Taksin Panpuy, Chayapa Phasomsap, Boonjit Deeaium, Pattiya Jootakarn.

Bamrasnaradura Infectious Diseases Institute, Nonthaburi, Thailand; Jurai Wongsawat, Rujanee Sunthornkachit, Visal Moolasart, Natawan Siripongpreeda, Supeda Thongyen, Piyawadee Chathaisong, Vilaiwan Prommool, Duangmanee Suwannamass, Simakan Waradejwinyoo, Nareopak Boonyarittipat, Thaniya Chiewcharn, Sirirat Likanonsakul, Chatiya Athichathana, Boonchuay Eampokalap, Wattana Sanchiem.

Srinagarind Hospital, Khon Kaen University, Khon Kaen, Thailand; Srinagarind Hospital, Khon Kaen University, Khon Kaen, Thailand; Pope Kosalaraksa, Pagakrong Lumbiganon, Chulapan Engchanil, Piangjit Tharnprisan, Chanasda Sopharak,Viraphong Lulitanond, Samrit Khahmahpahte, Ratthanant Kaewmart, Prajuab Chaimanee, Mathurot Sala, Thaniita Udompanit,Ratchadaporn Wisai,Somjai Rattanamanee, Yingrit Chantarasuk, Sompong Sarvok, Yotsombat Changtrakun, Soontorn Kunhasura, Sudthanom Kamollert.

Queen Savang Vadhana Memorial Hospital, Chonburi, Thailand; Wicharn Luesomboon, Dr. Pairuch Eiamapichart, Dr. Tanate Jadwattanakul, Isara Limpetngam, Daovadee Naraporn, Pornpen Mathajittiphun, Chatchadha Sirimaskul, Woranun Klaihong, Pipat Sittisak, Tippawan Wongwian, Kansiri Charoenthammachoke, Pornchai Yodpo.

Nakornping Hospital, Chiang Mai, Thailand; Suparat Kanjanavanit, Maneerat Ananthanavanich, Penpak Sornchai, Thida Namwong, Duangrat Chutima, Suchitra Tangmankhongworakun, Pacharaporn Yingyong, Juree Kasinrerk, Montanee Raksasang, Pimporn Kongdong, Siripim Khampangkome, Suphanphilat Thong-Ngao, Sangwan Paengta, Kasinee Junsom, Ruttana Khuankaew M, Parichat Moolsombat, Duanpen Khuttiwung, Chanannat Chanrin.

Chiangrai Regional Hospital, ChiangRai, Thailand; Rawiwan Hansudewechakul, Yaowalak Jariyapongpaiboon, Chulapong Chanta, Areerat Khonponoi, Chaniporn Yodsuwan, Warunee Srisuk, Pojjavitt Ussawawuthipong, Yupawan Thaweesombat, Polawat Tongsuk, Chaiporn Kumluang, Ruengrit Jinasen, Noodchanee Maneerat, Kajorndej Surapanichadul, Pornpinit Donkaew.

Prapokklao Hospital, Chantaburi,Thailand; Chaiwat Ngampiyaskul, Naowarat Srisawat, Wanna Chamjamrat, Sayamol Wattanayothin, Pornphan Prasertphan, Tanyamon Wongcheeree, Pisut Greetanukroh, Chataporn Imubumroong, Pathanee Teirsonsern.

Research Institute for Health Sciences, Chiang Mai University, Chiang Mai, Thailand; Virat Sirisanthana, Linda Aurpibul, Pannee Visrutaratna, Siriporn Taphey, Tawalchaya Cholecharoentanan, Nongyow Wongnum, Chintana Khamrong, Rassamee Kaewvichit, Kittipong Rungroengthanakit.

Chiang Mai University: Kulvadee Thongpibul

## Disclaimer

The content of this manuscript is solely the responsibility of the authors and does not necessarily represent the official views of any of the institutions mentioned above, the U.S. Department of the Army or the U.S. Department of Defense, the National Institutes of Health, the Department of Health and Human Services, or the United States government, nor does mention of trade names, commercial products, or organizations imply endorsement by the Thai Red Cross AIDS Research Centre.

