## supplement for "Predicting neurodevelopmental outcomes in children with perinatal HIV using a novel machine learning algorithm"

**SUPPLEMENTARY MATERIAL**

**Feature 1. Cluster of Mean Corpuscular Hemoglobin, CBCL Affective Problems Score, and total CBCL Score**

**Figure S1. Features 1-25.** Discrimination plots for the top 25 feature/feature clusters that predicted neurocognitive outcomes in children with pHIV. Feature plots are presented in order of predictive importance.

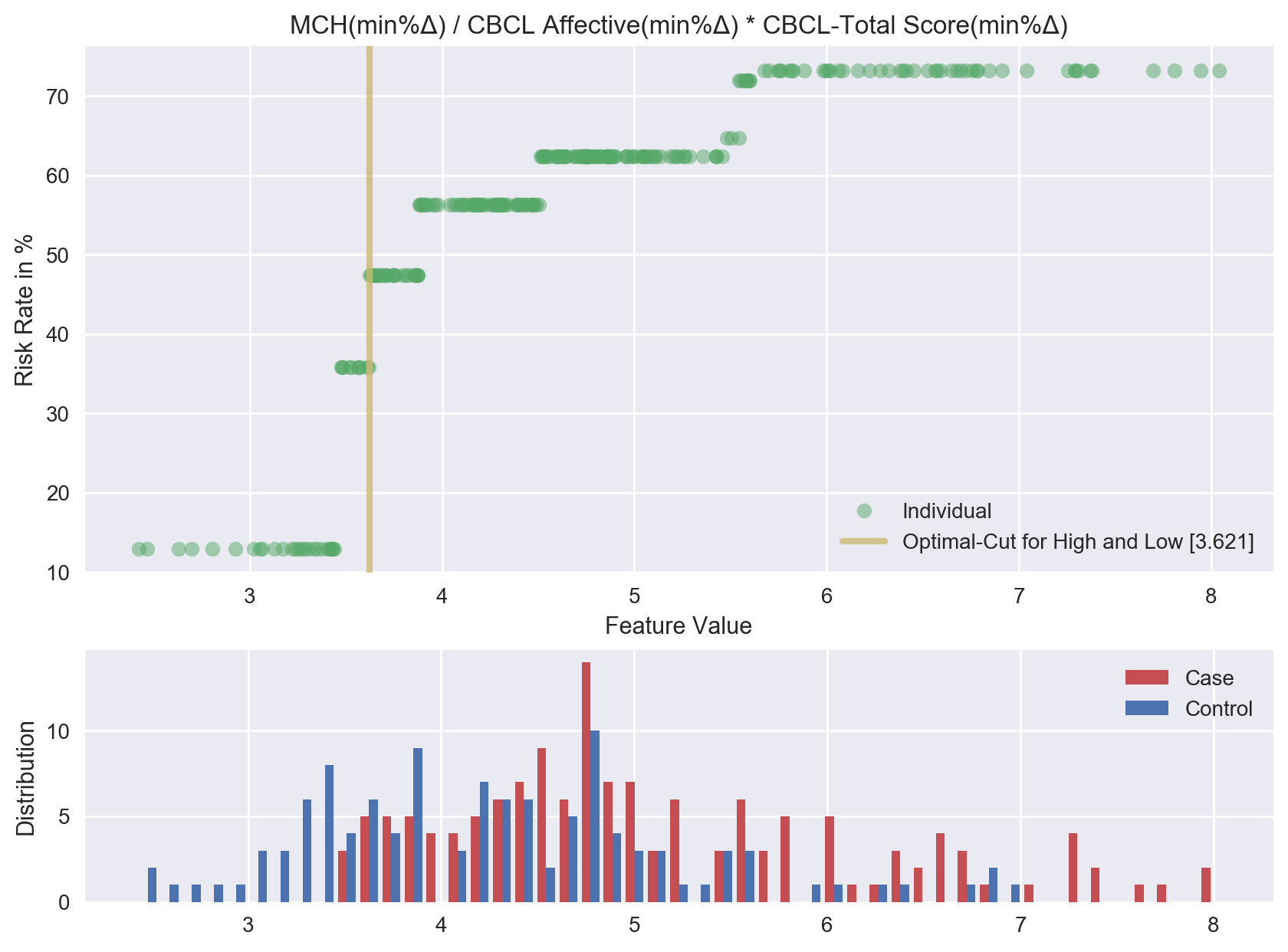

**Feature 2. Cluster of Triglycerides, CBCL Affective Problems Score, and CBCL Anxious Subscale**

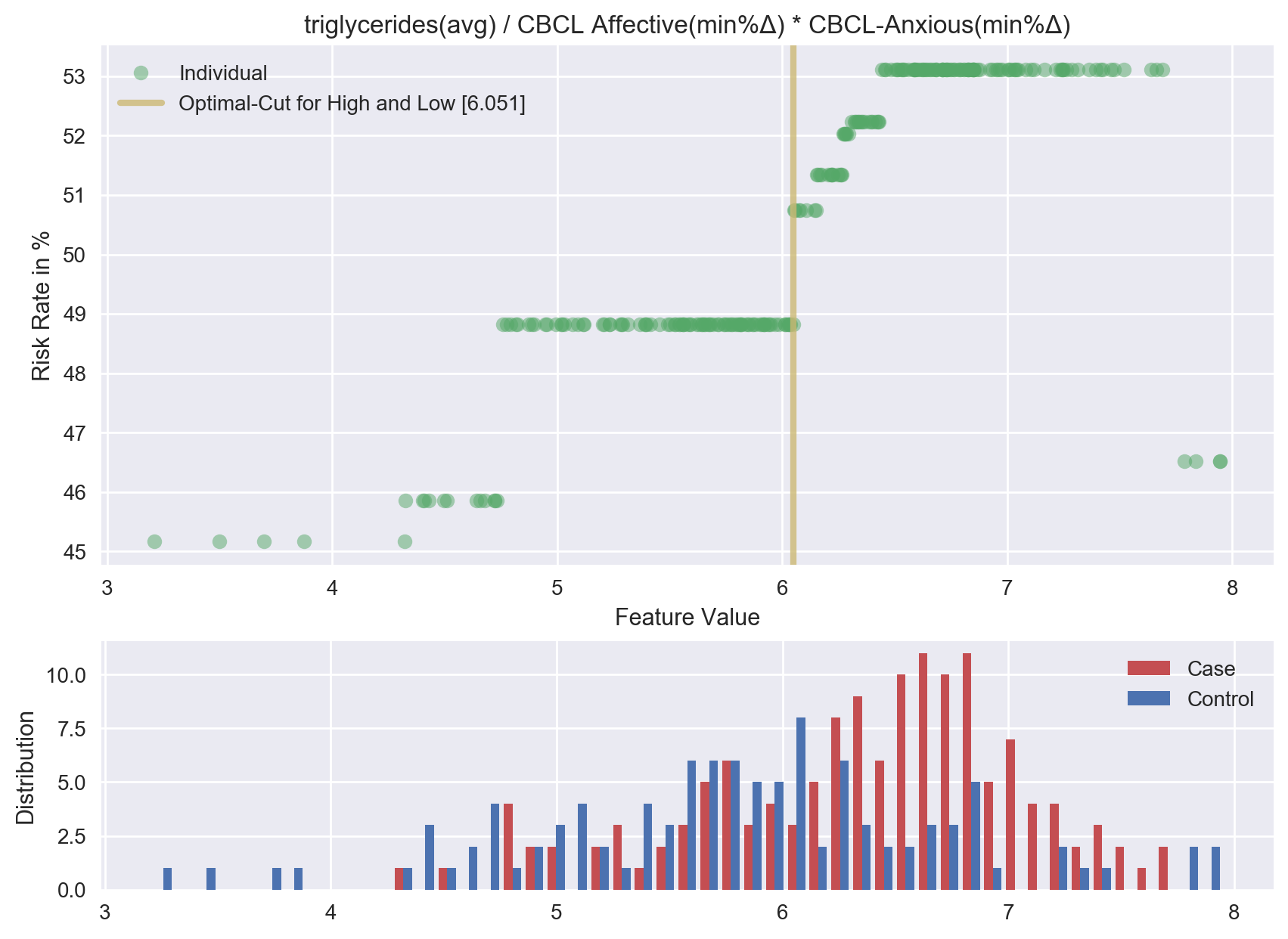

**Feature 3. Lymphocytes**

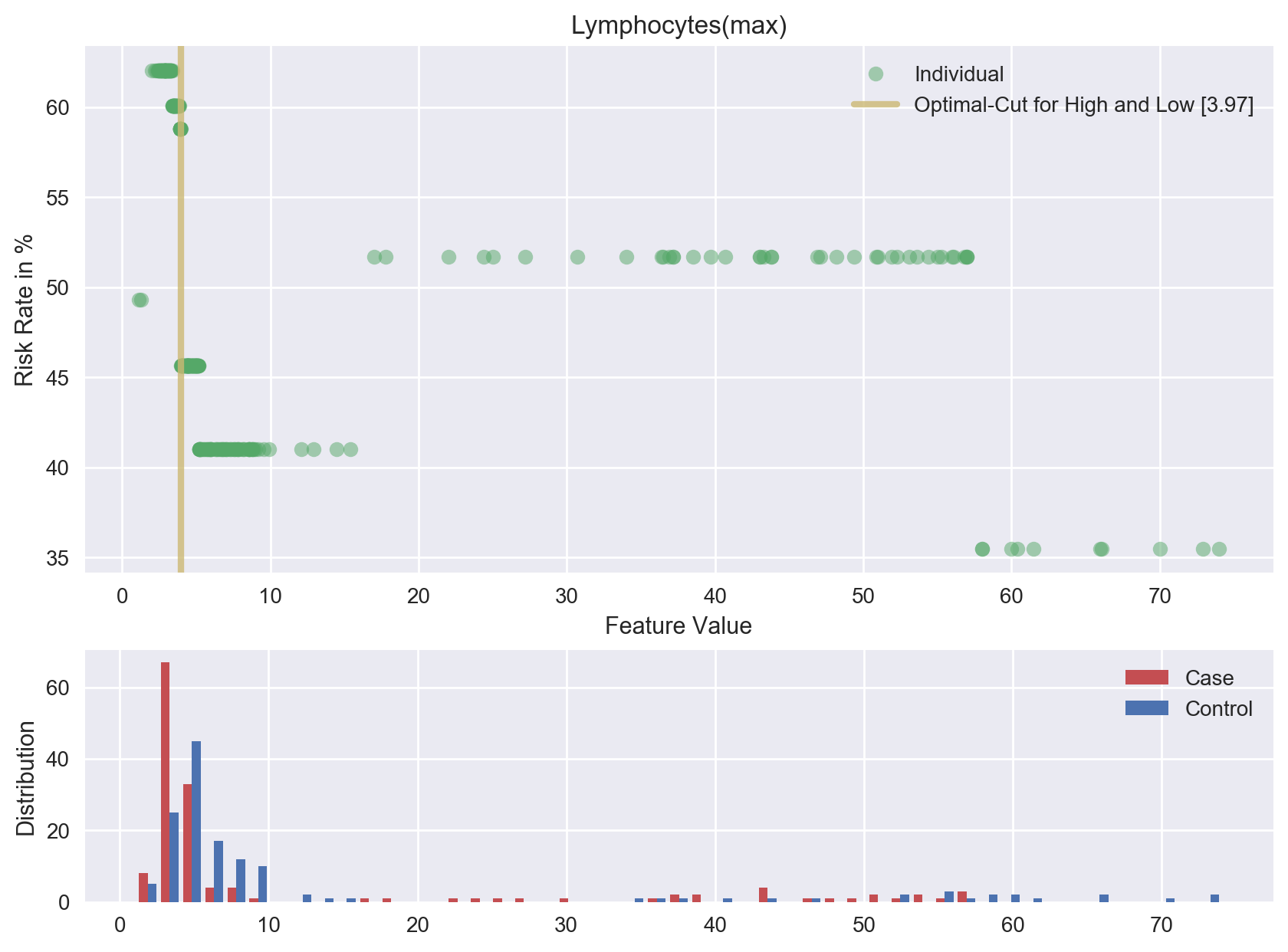

**Feature 4. CD8 Cell Count**

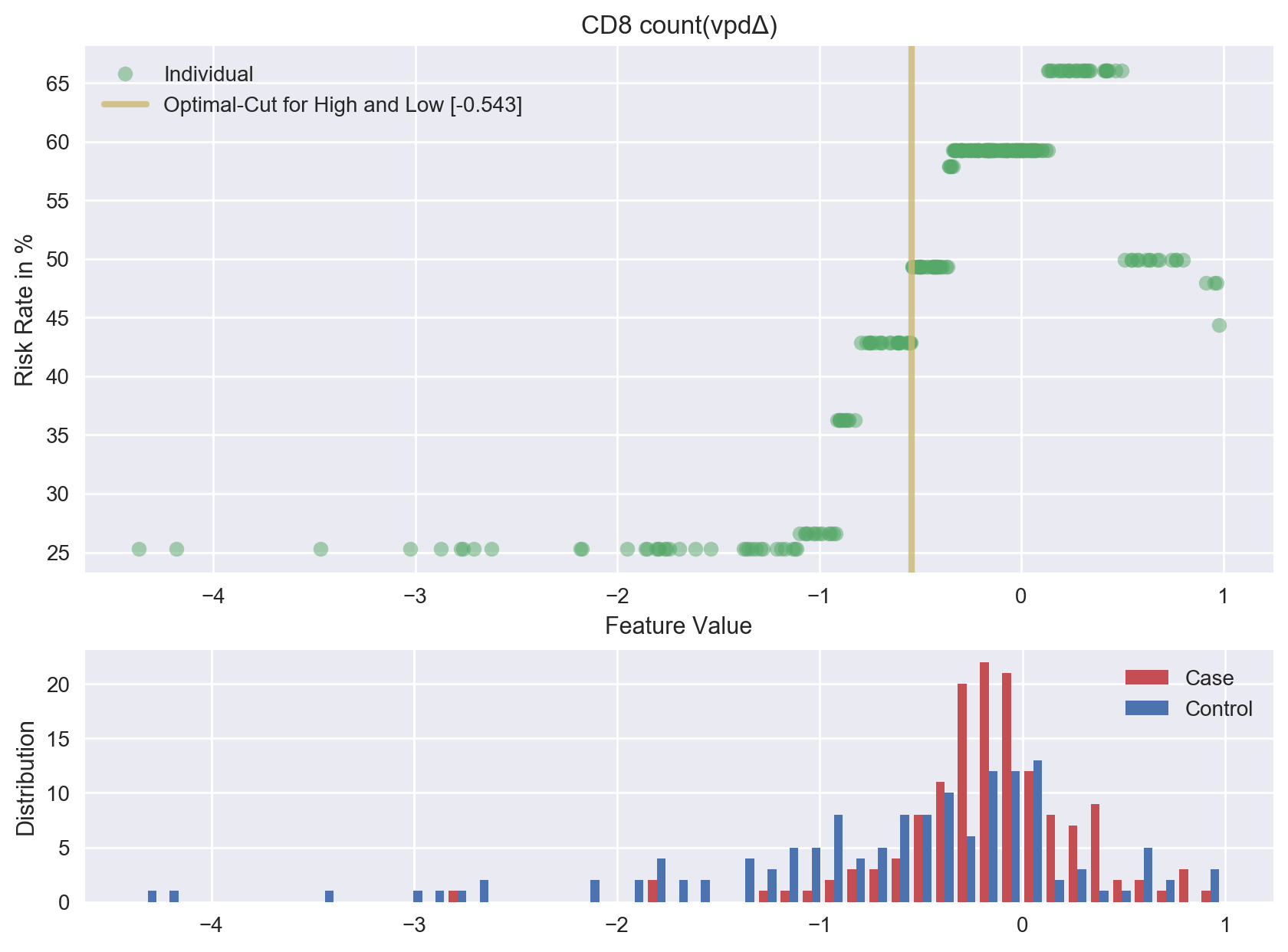

**Feature 5. Cluster of CBCL Anxious Subscale, CBCL Somatic Subscale, and total CBCL Score**

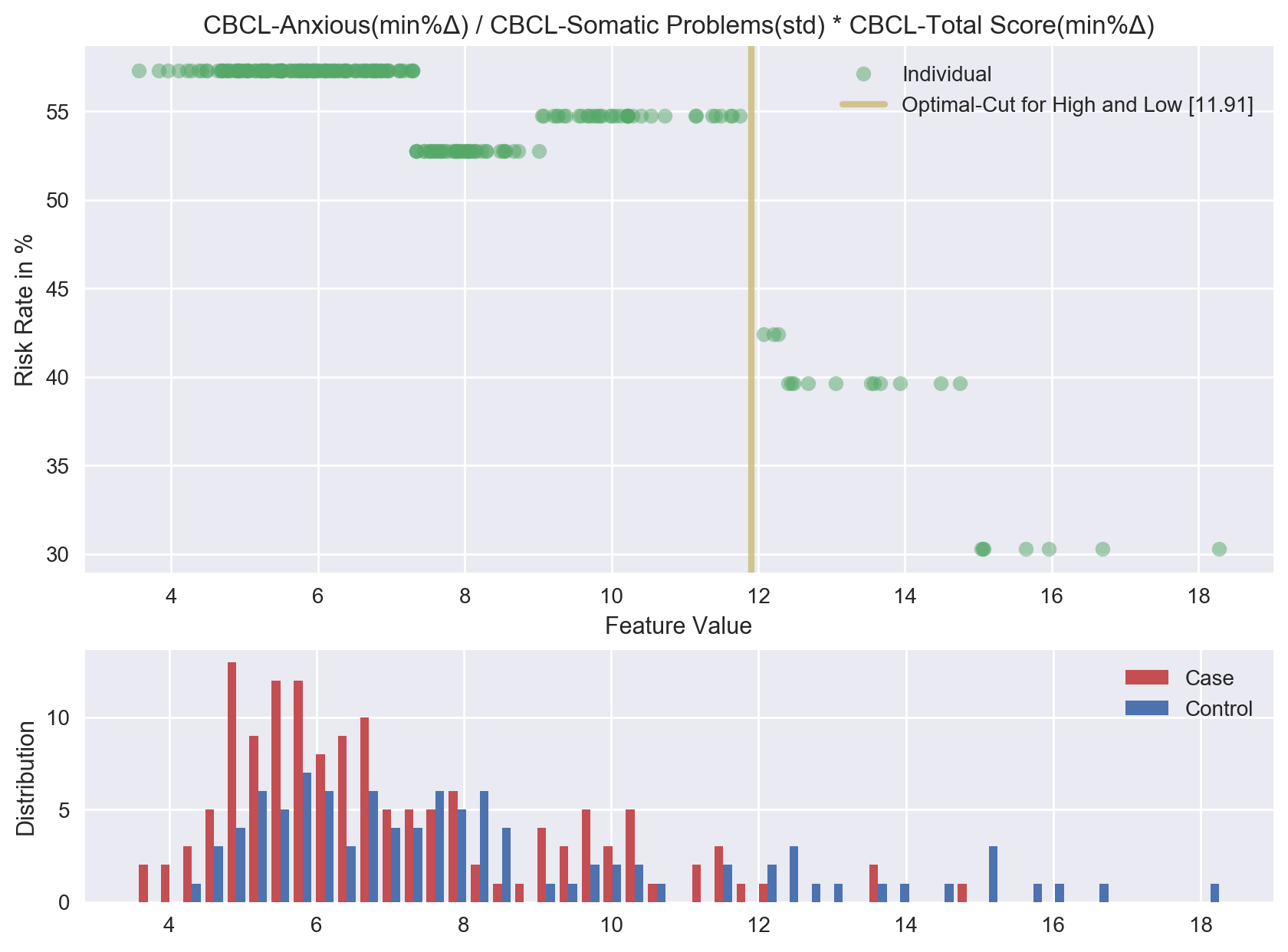

**Feature 6. Mean Corpuscular Hemoglobin**

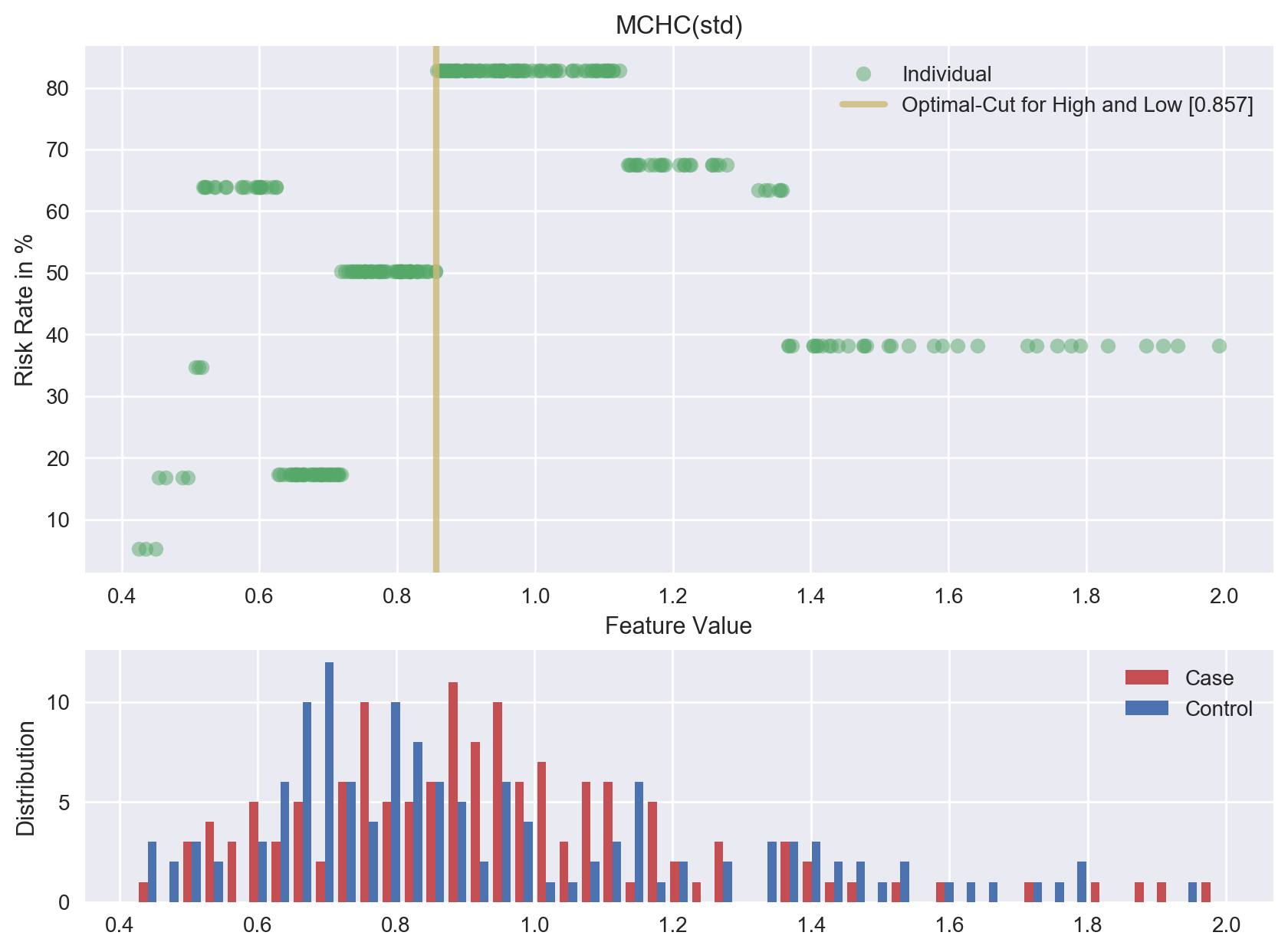

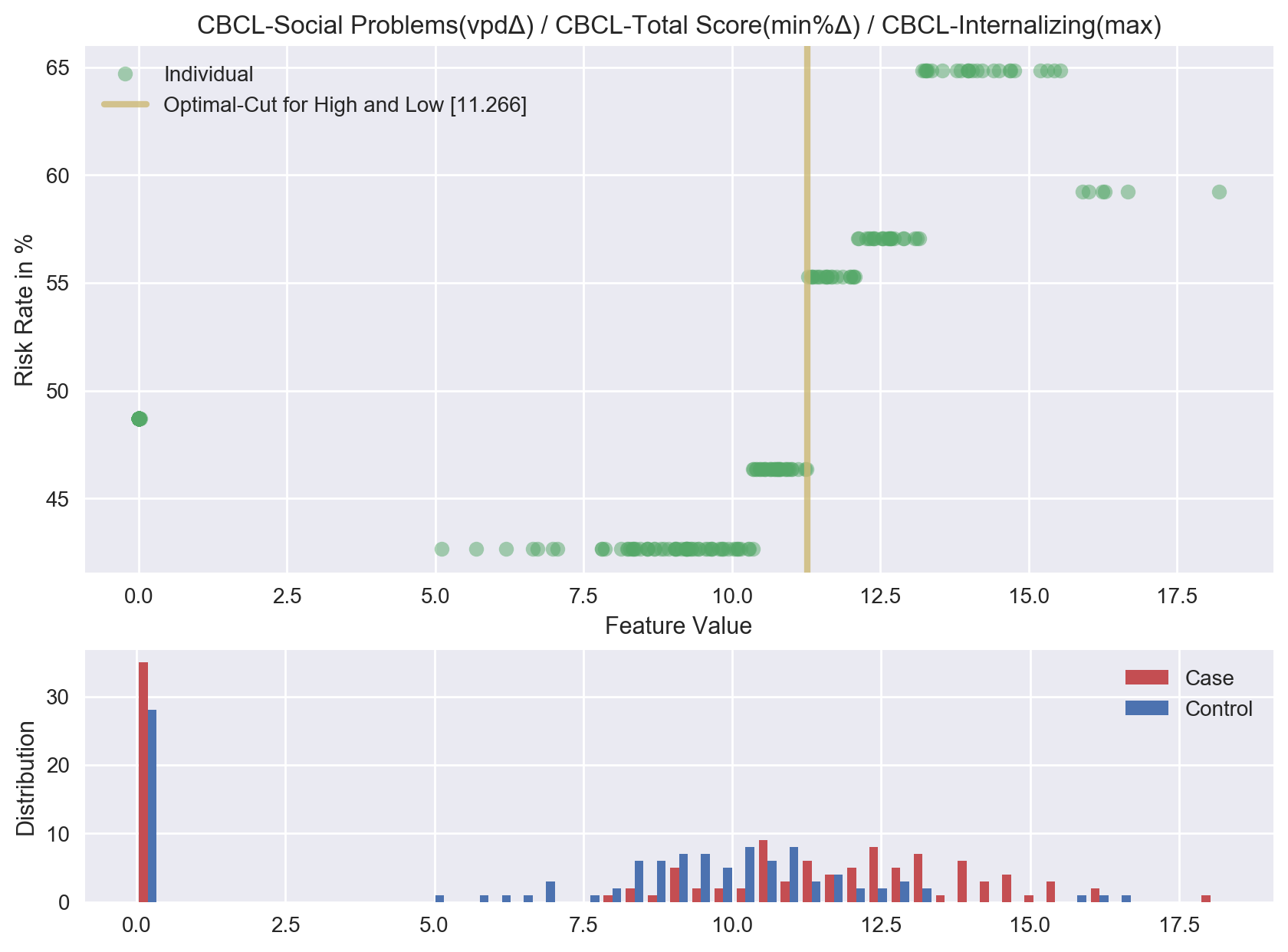

**Feature 7. Cluster of CBCL Social Problems Subscale , total CBCL Score, and CBCL Internalizing Score**

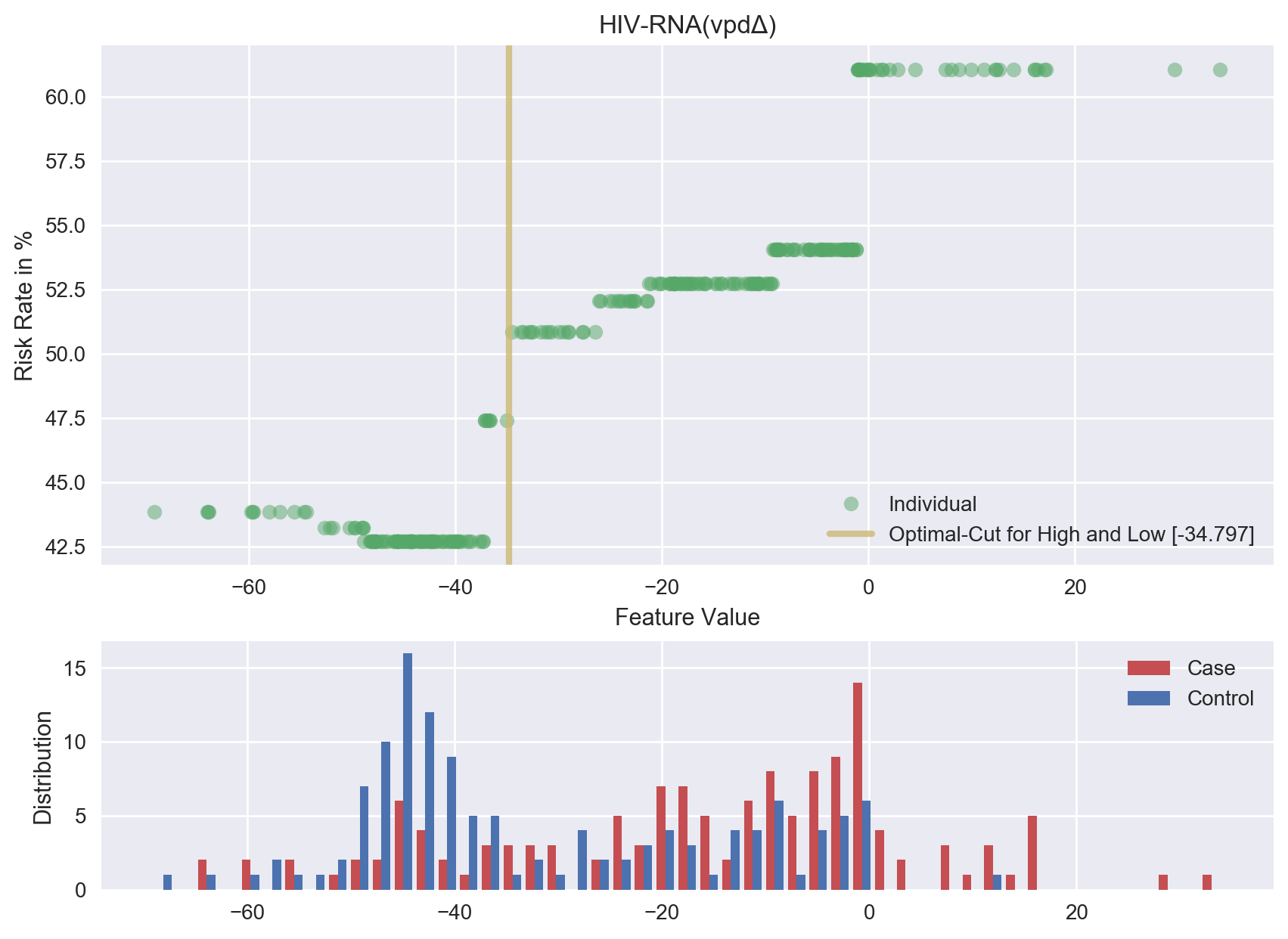

**Feature 8. HIV-RNA**

**Feature 9. Cluster of Red Blood Cell, CBCL Affective Problems Score, and CBCL Social Competence Scale Score**

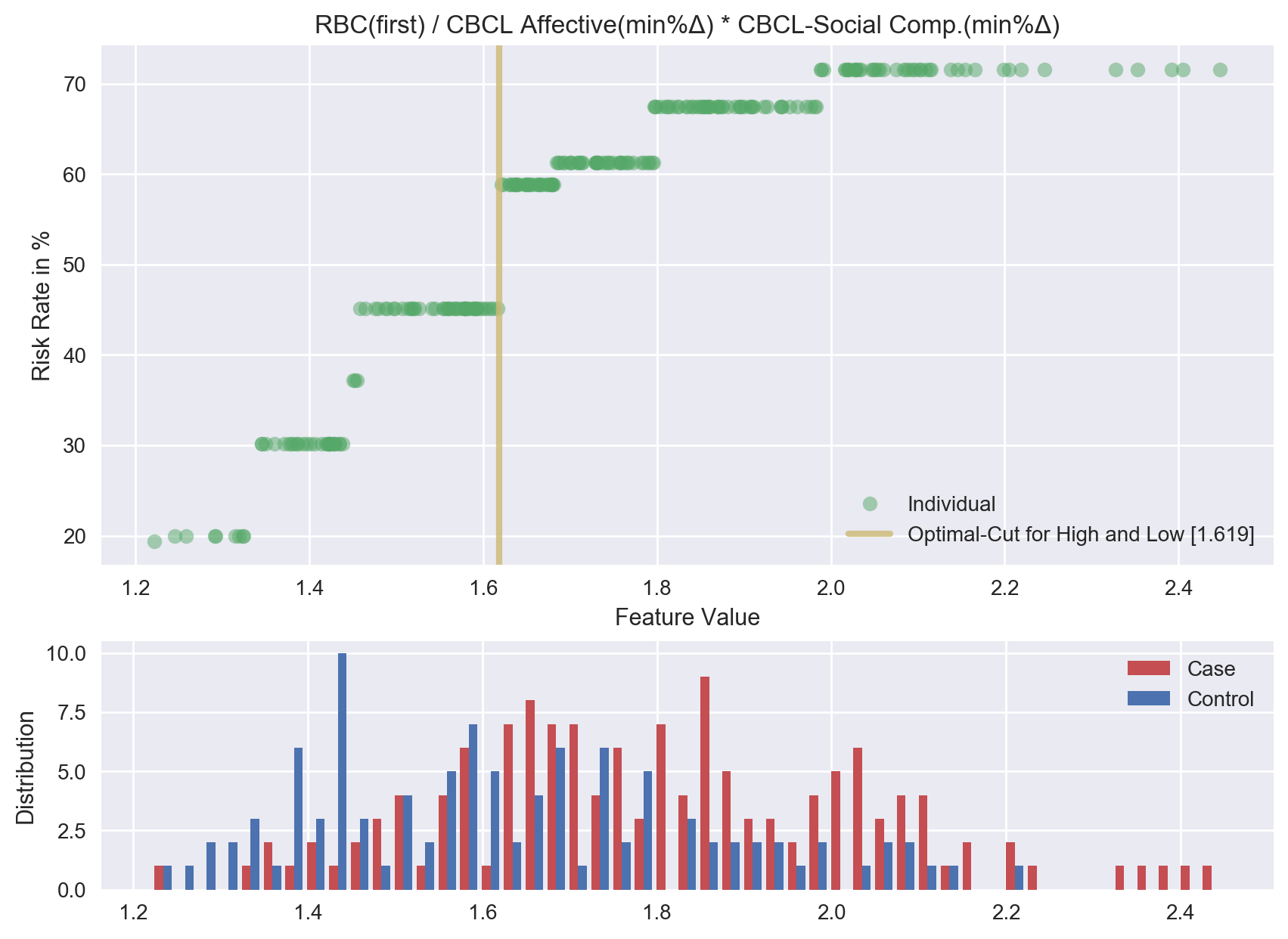

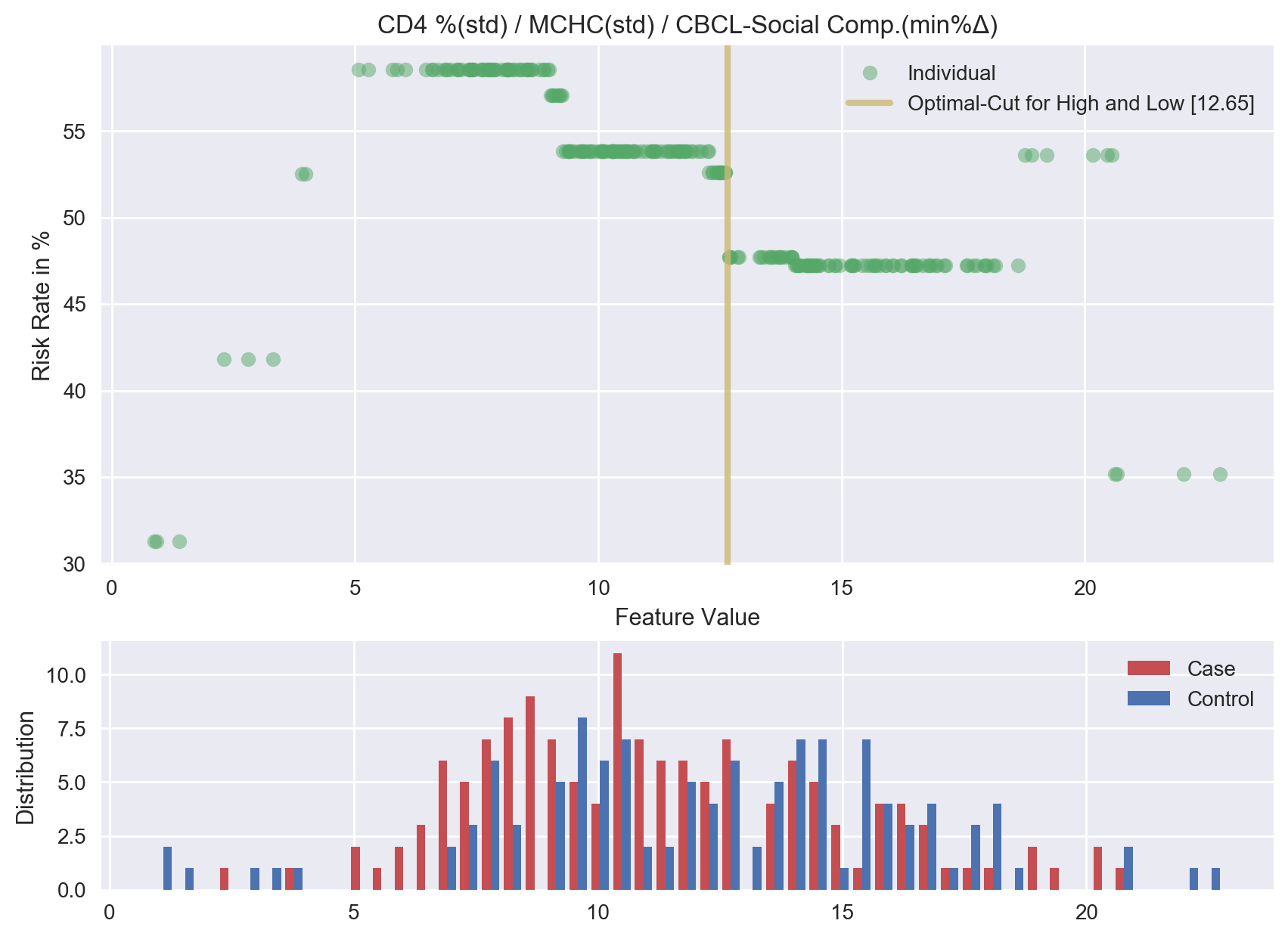

**Feature 10. Cluster of CD4 Percentage, Mean Corpuscular Hemoglobin Concentration, and CBCL Social Competence Scale Score**

**Feature 11. Cluster of Red Blood Cell, CD4 Percentage, and CBCL Anxious Subscale**

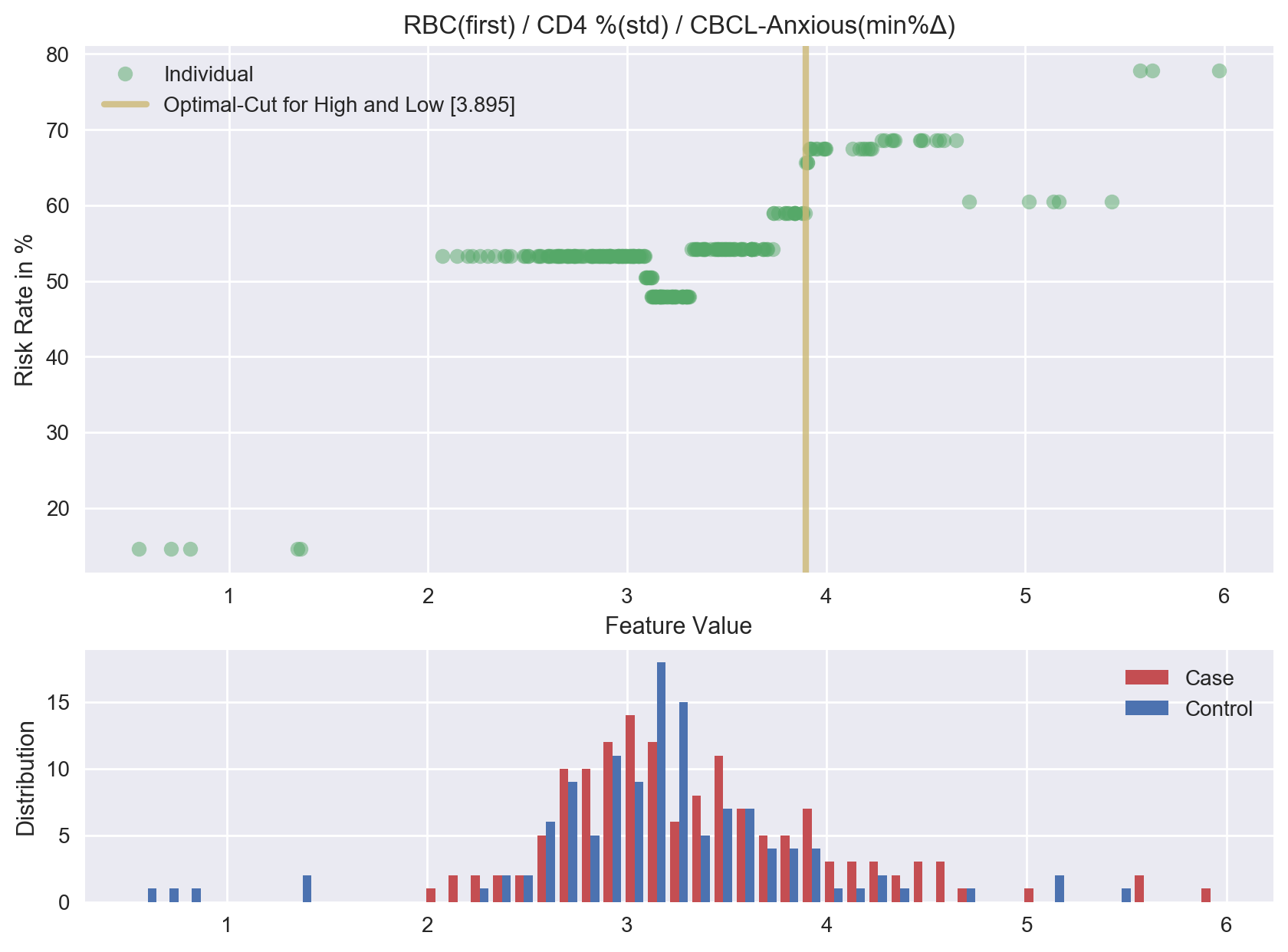

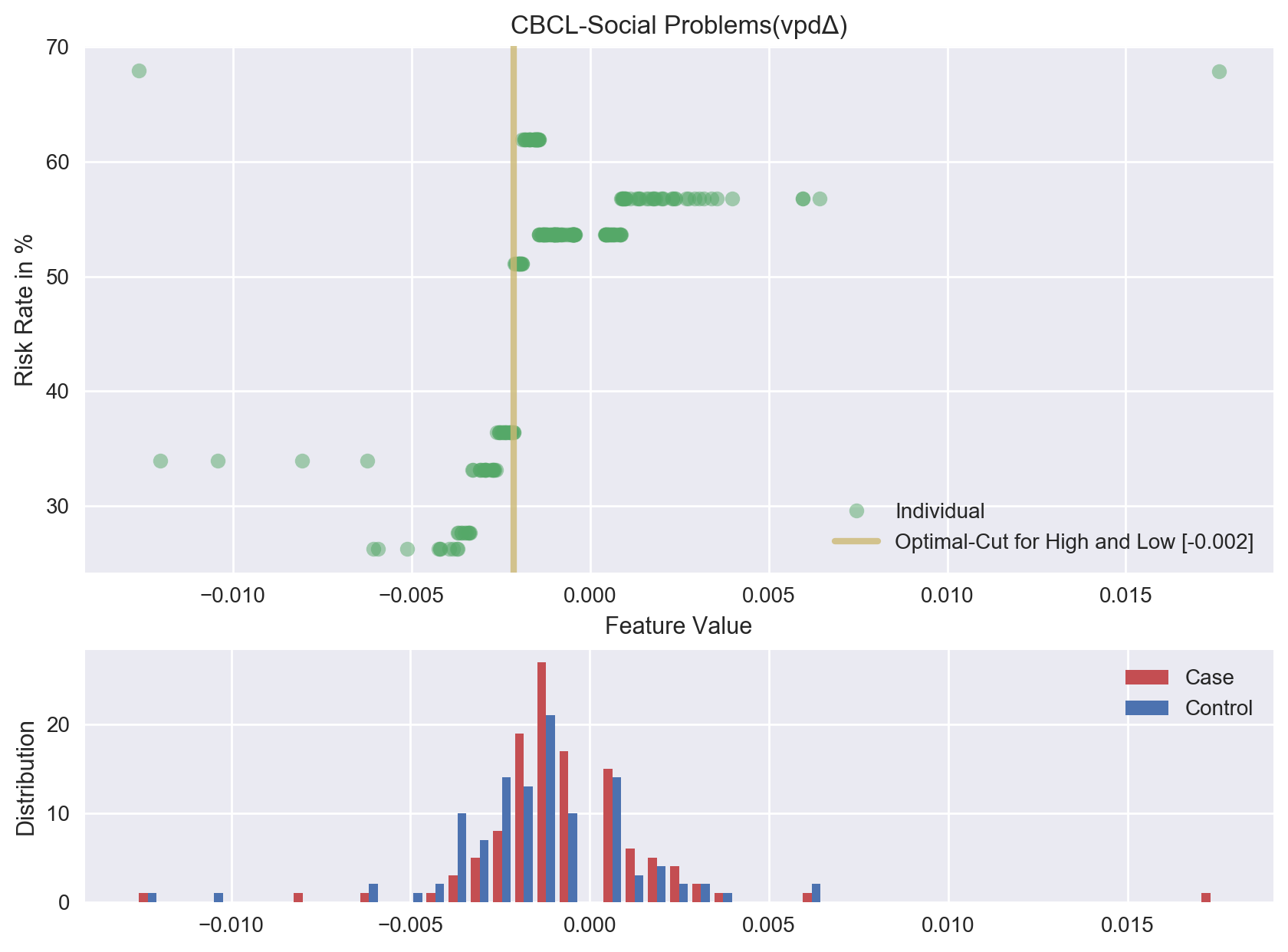

**Feature 12. CBCL Social Problems Subscale**

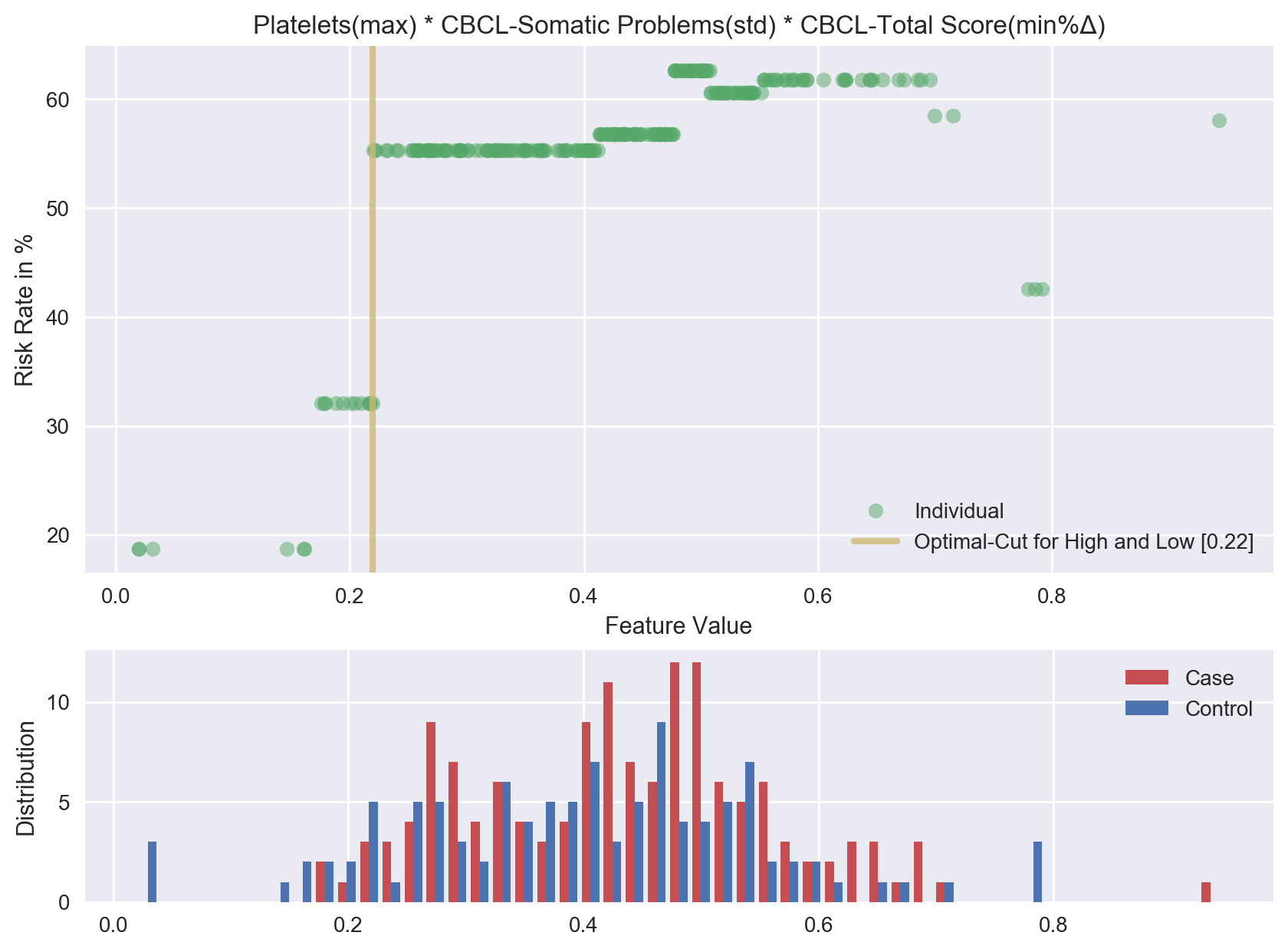

**Feature 13. Cluster of Platelets, CBCL Somatic Problems Score, and total CBCL Score**

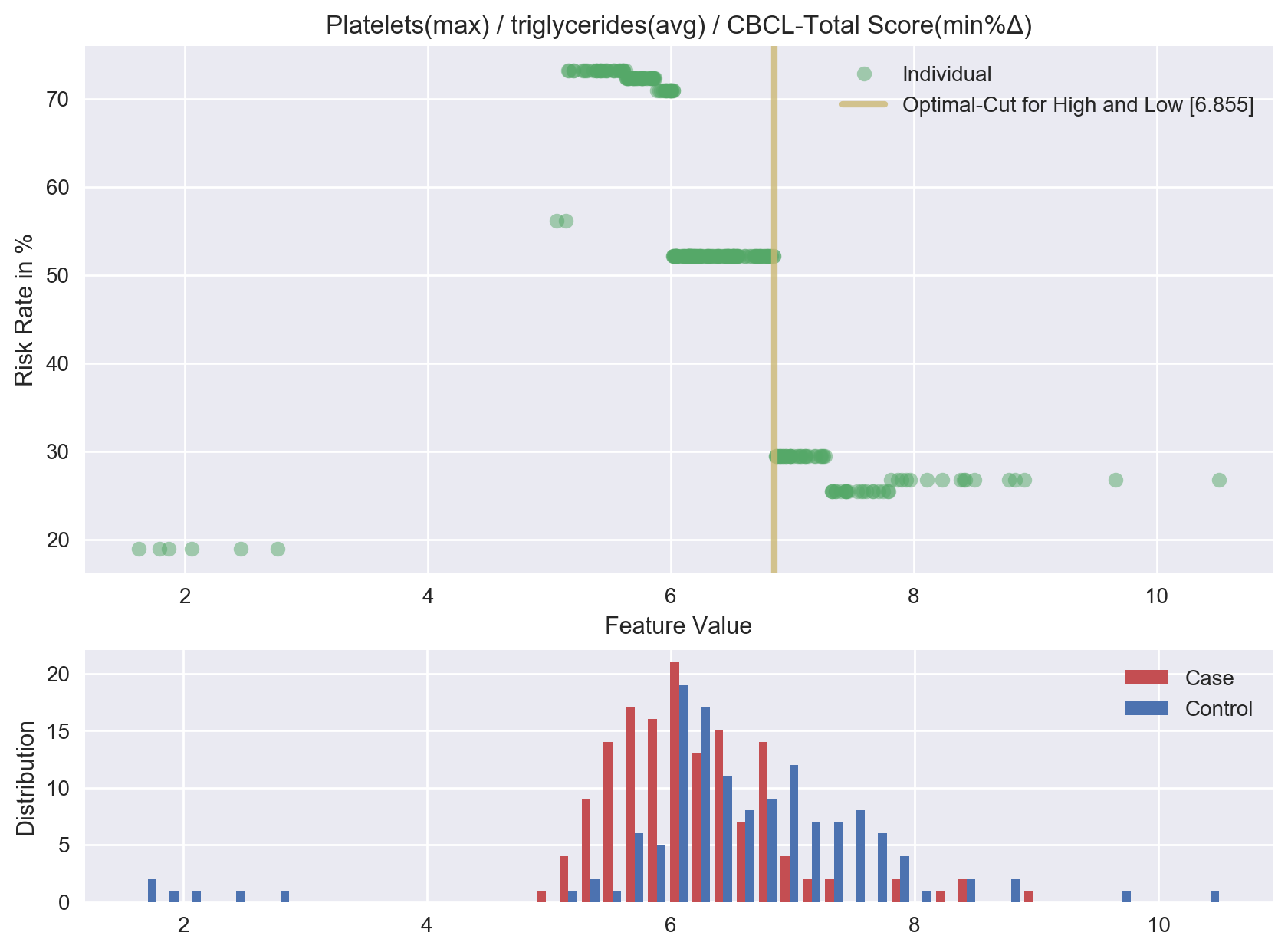

**Feature 14. Cluster of Platelets, Triglycerides, and total CBCL Score**

**Feature 15. Triglycerides**

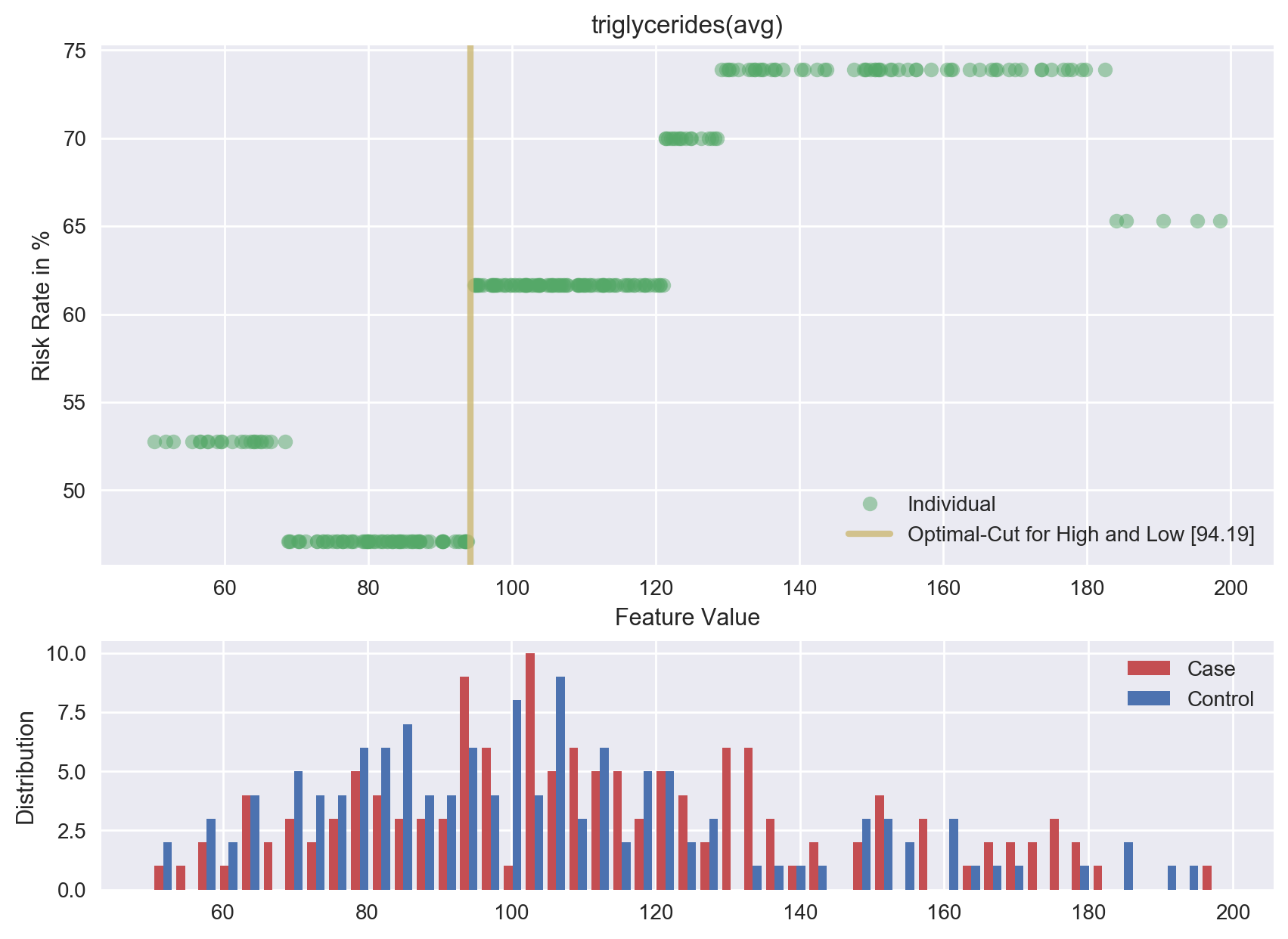

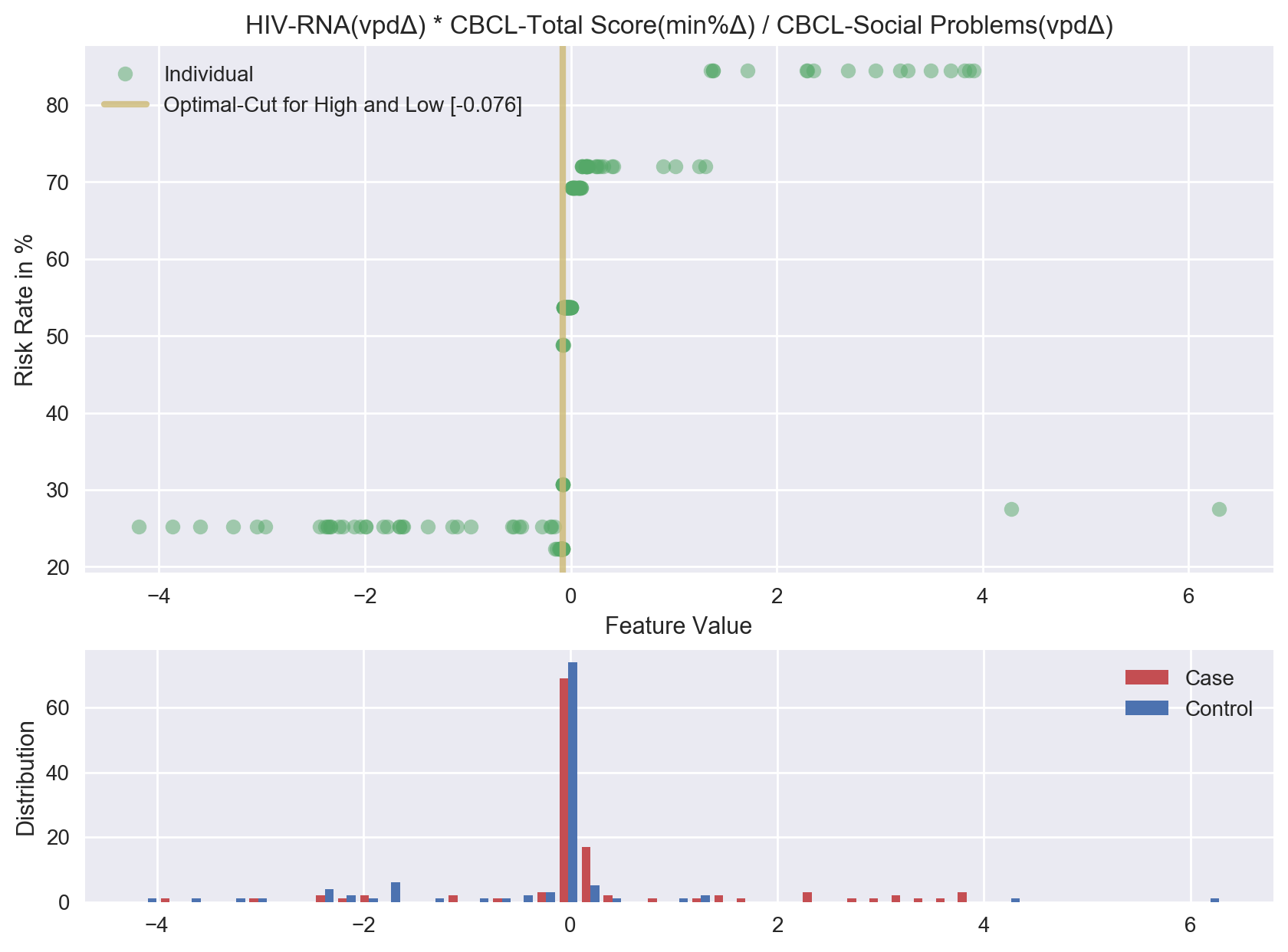

**Feature 16. Cluster of HIV-RNA, total CBCL Score, and CBCL Social Problems Subscale**

**Feature 17. Cluster of Mean Corpuscular Hemoglobin Concentration and CBCL Somatic Problems Score**

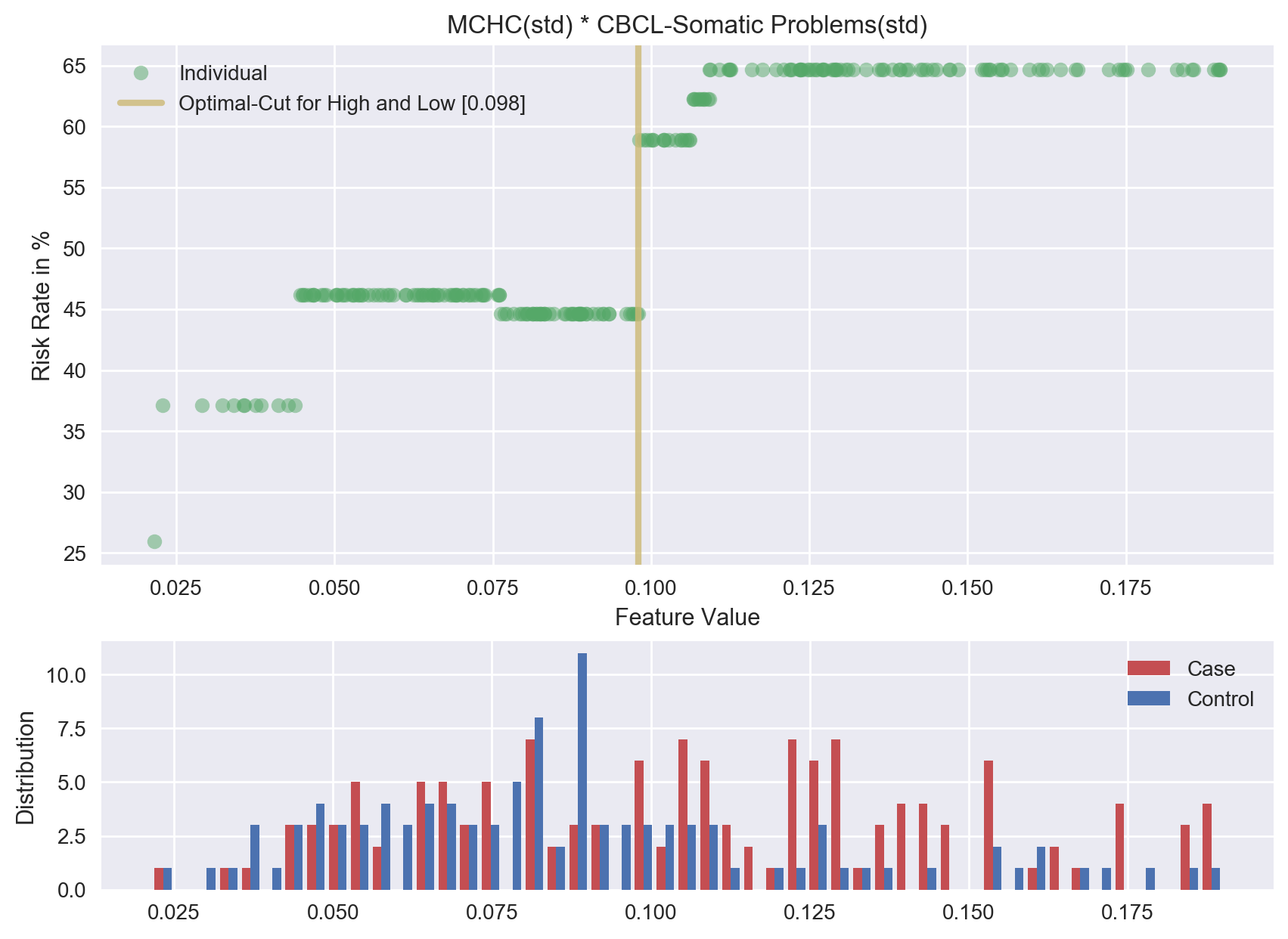

**Feature 18. Cluster of Triglycerides and Platelets**

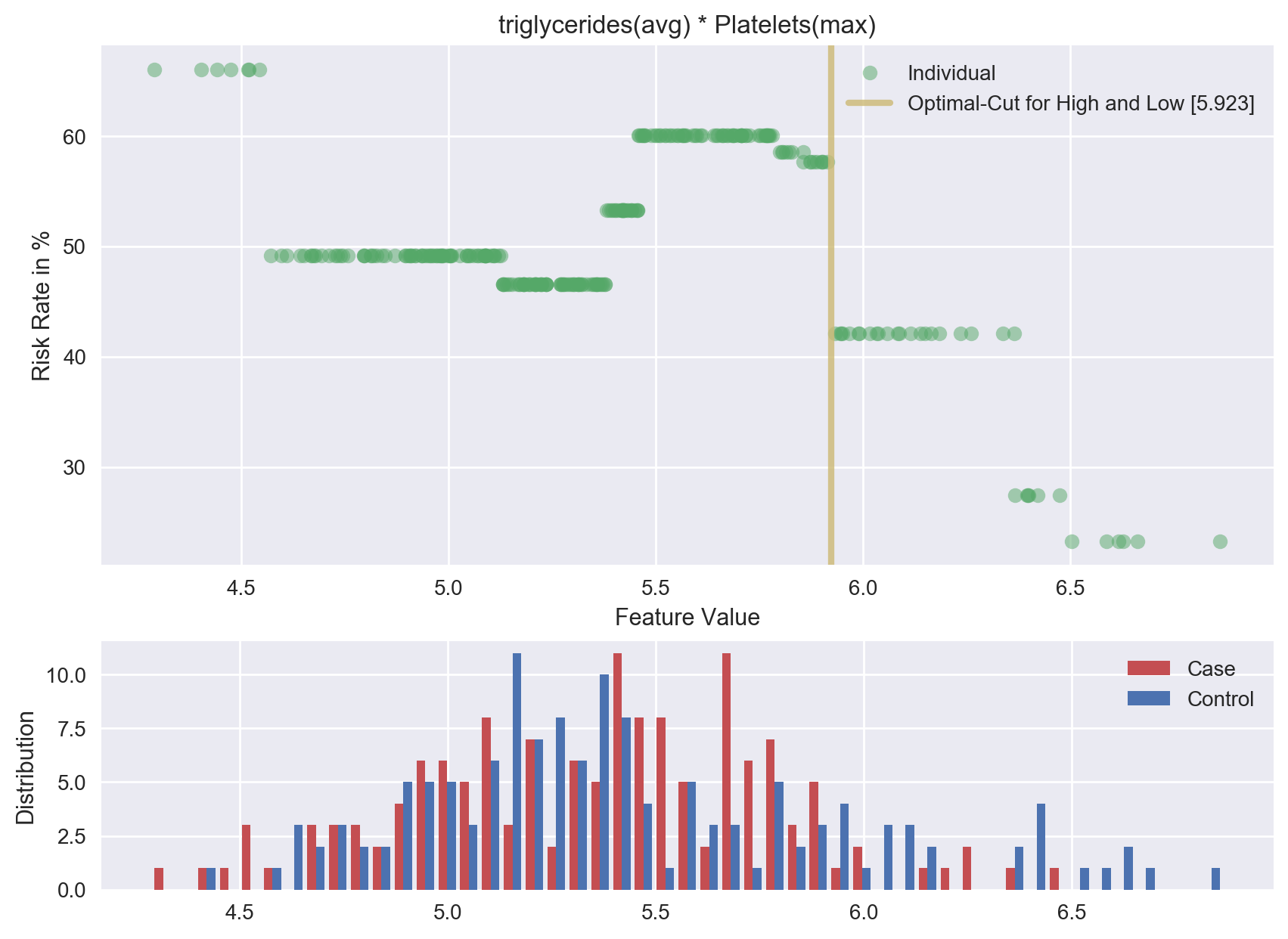

**Feature 19. CD4 Percentage**

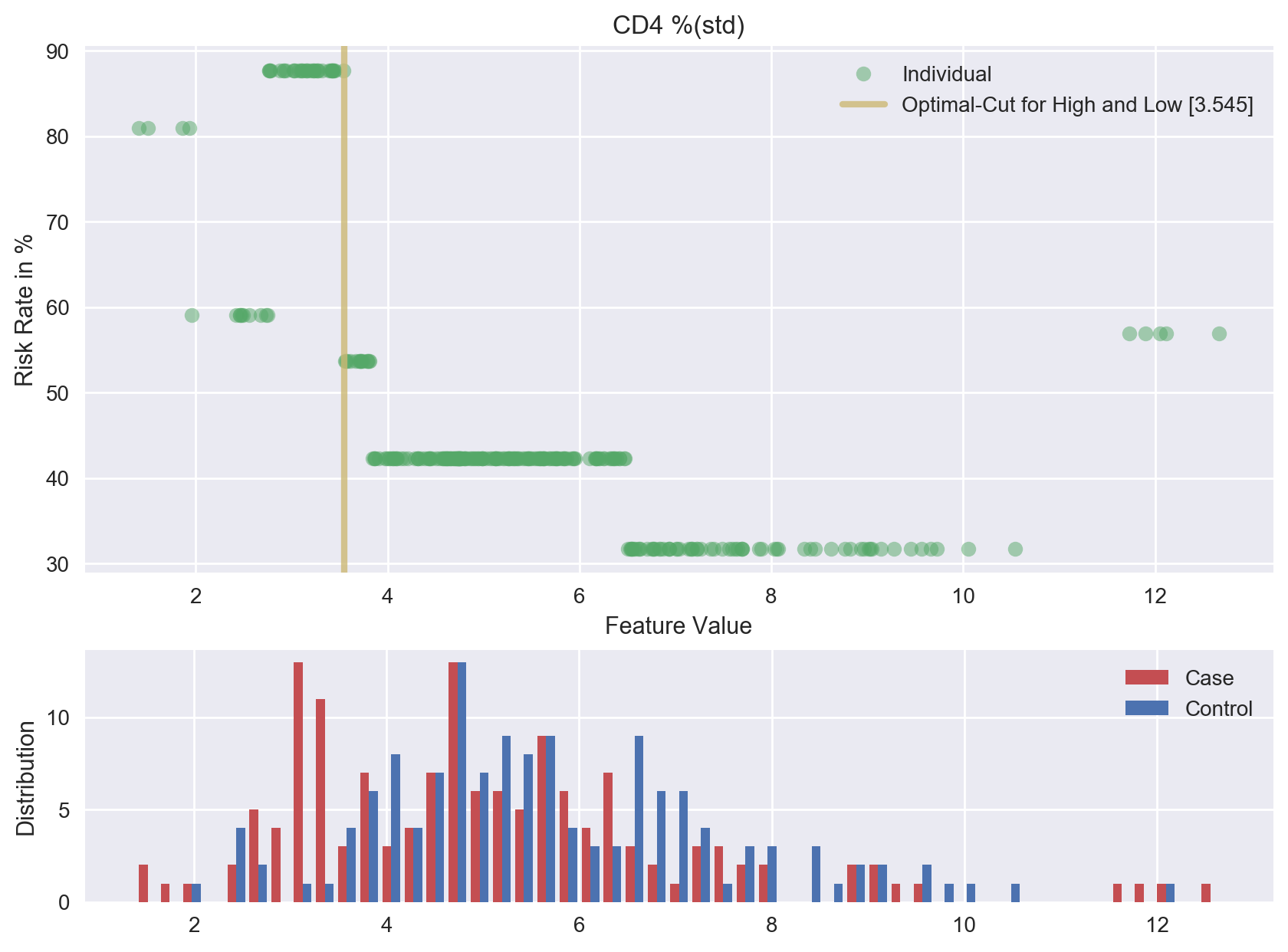

**Feature 20. Cluster of HIV-RNA, CBCL Affective Problems Score, and CBCL Activities Competence Scale Score**

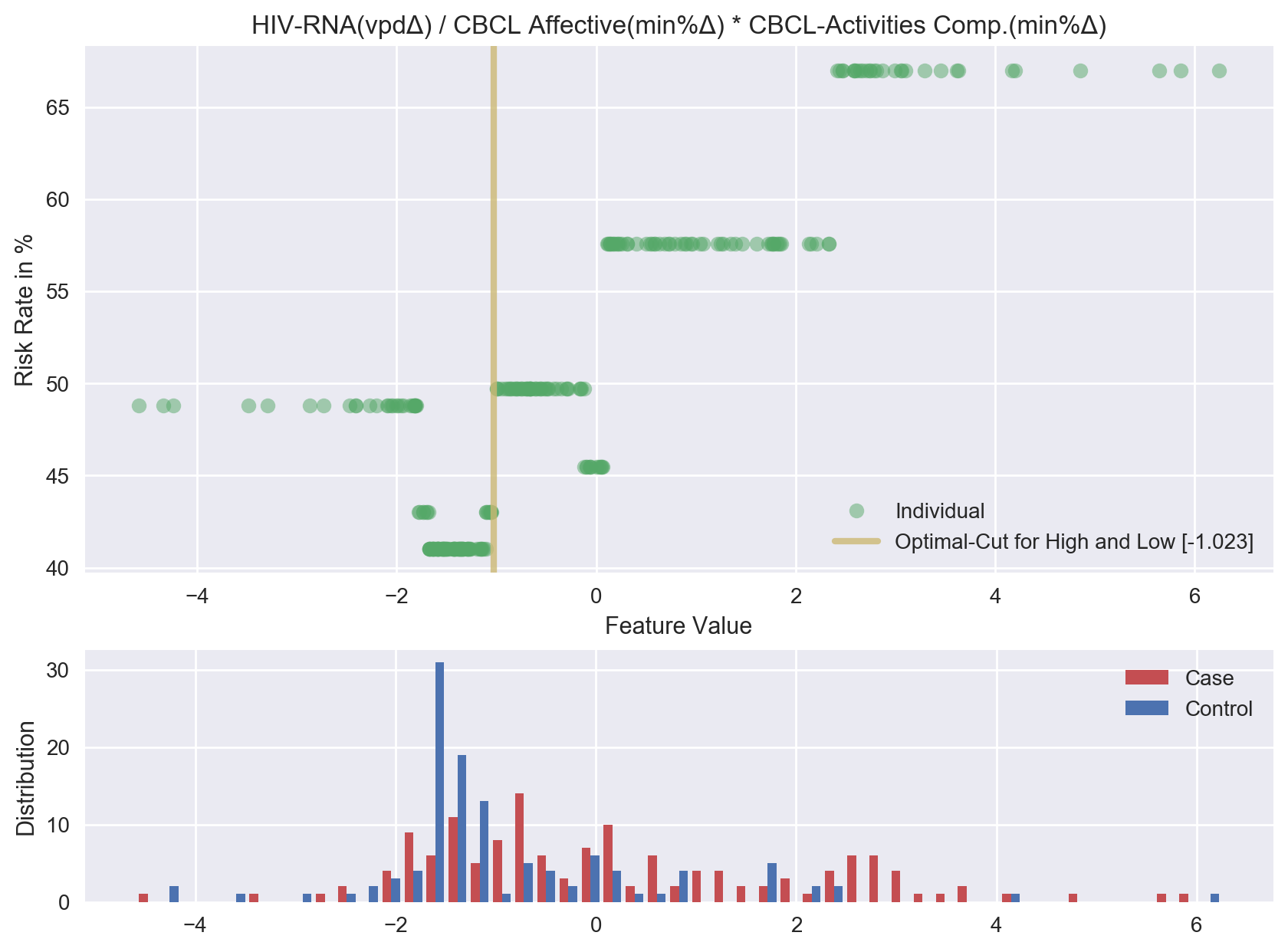

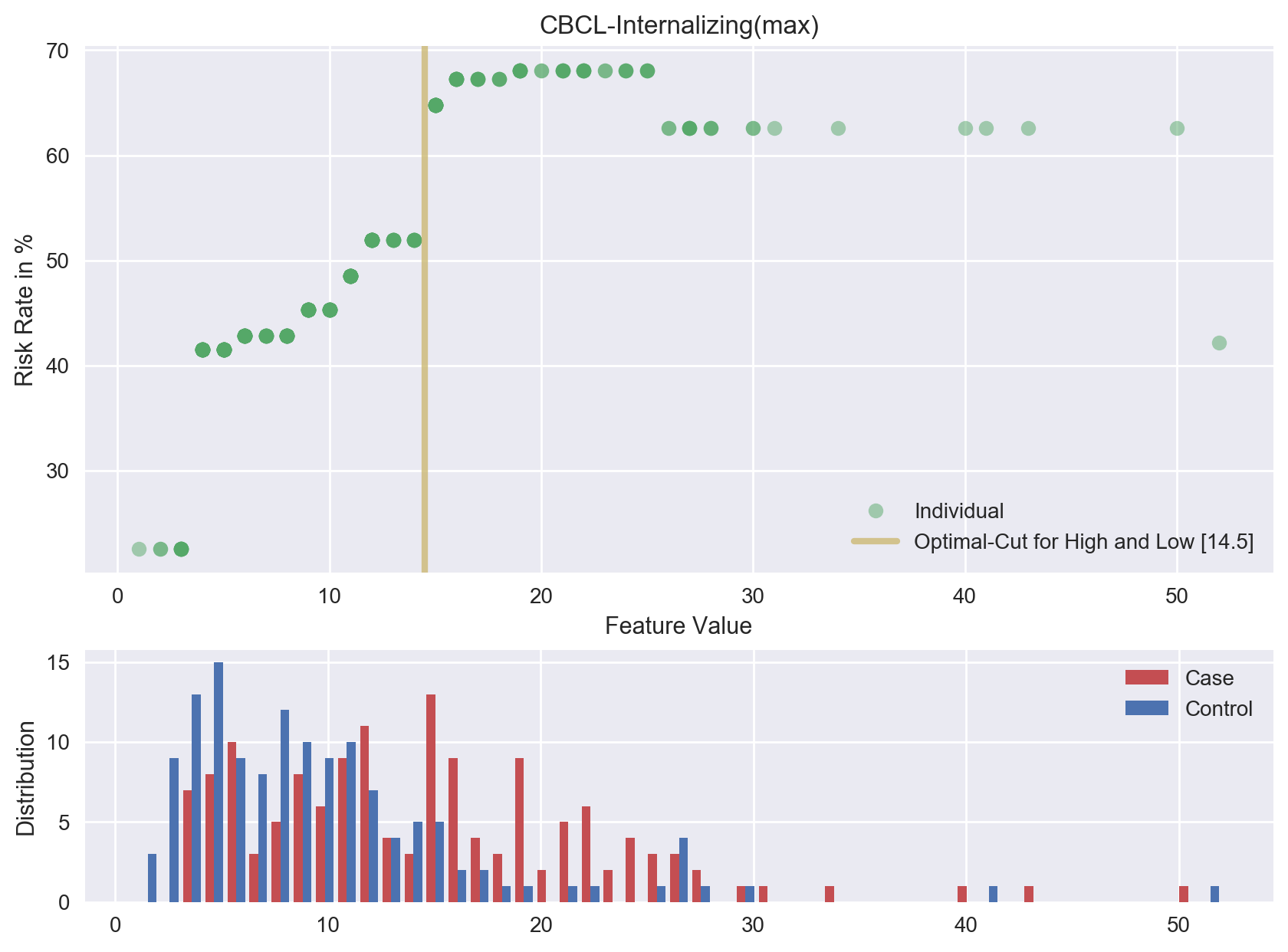

**Feature 21. CBCL Internalizing Subscale**

**Feature 22. Cluster of Triglycerides and CBCL Affective Problems Score**

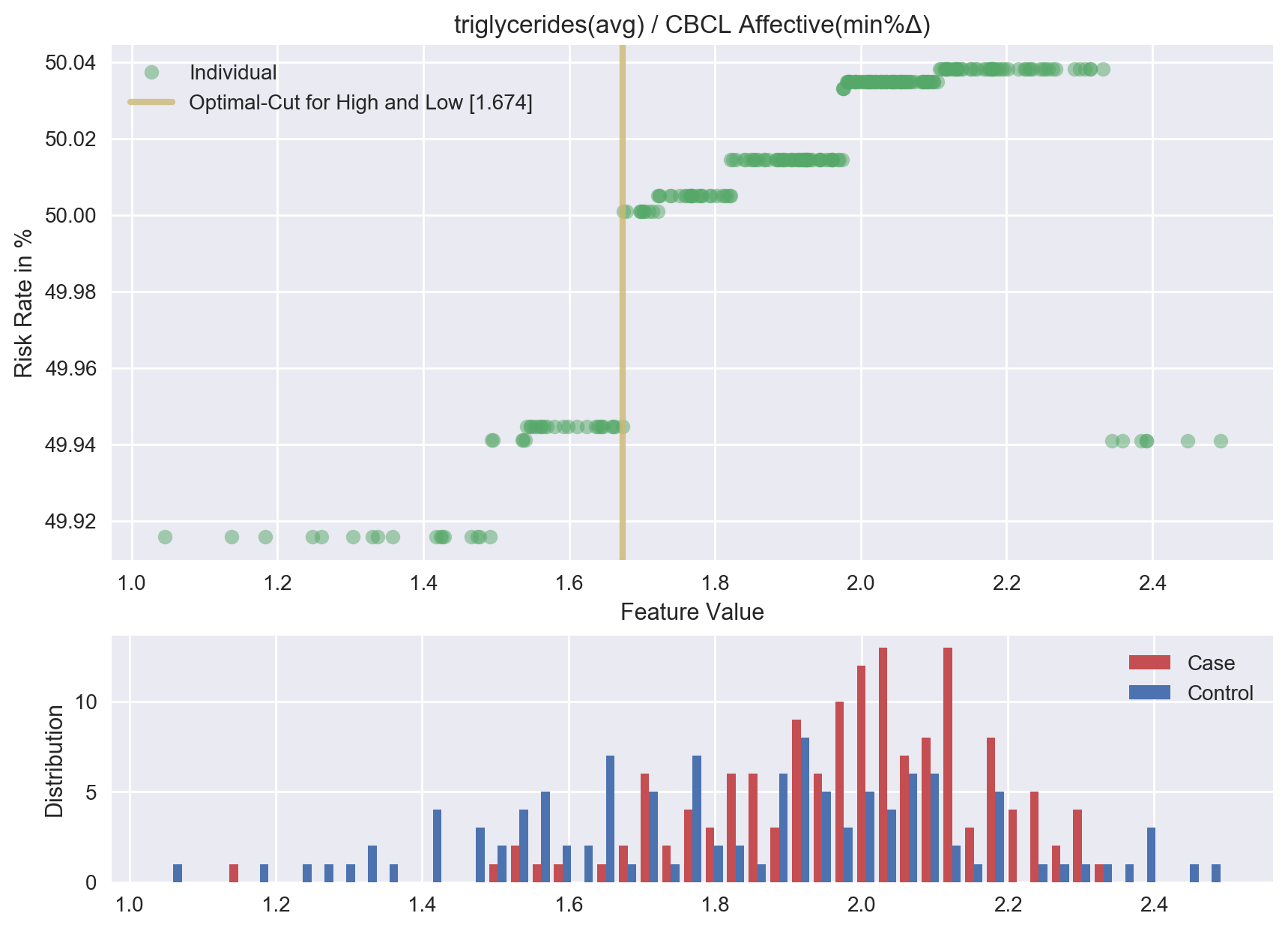

**Feature 23. Cluster of CD8 Cell Count, CBCL Social Problems Subscale, and HIV-RNA**

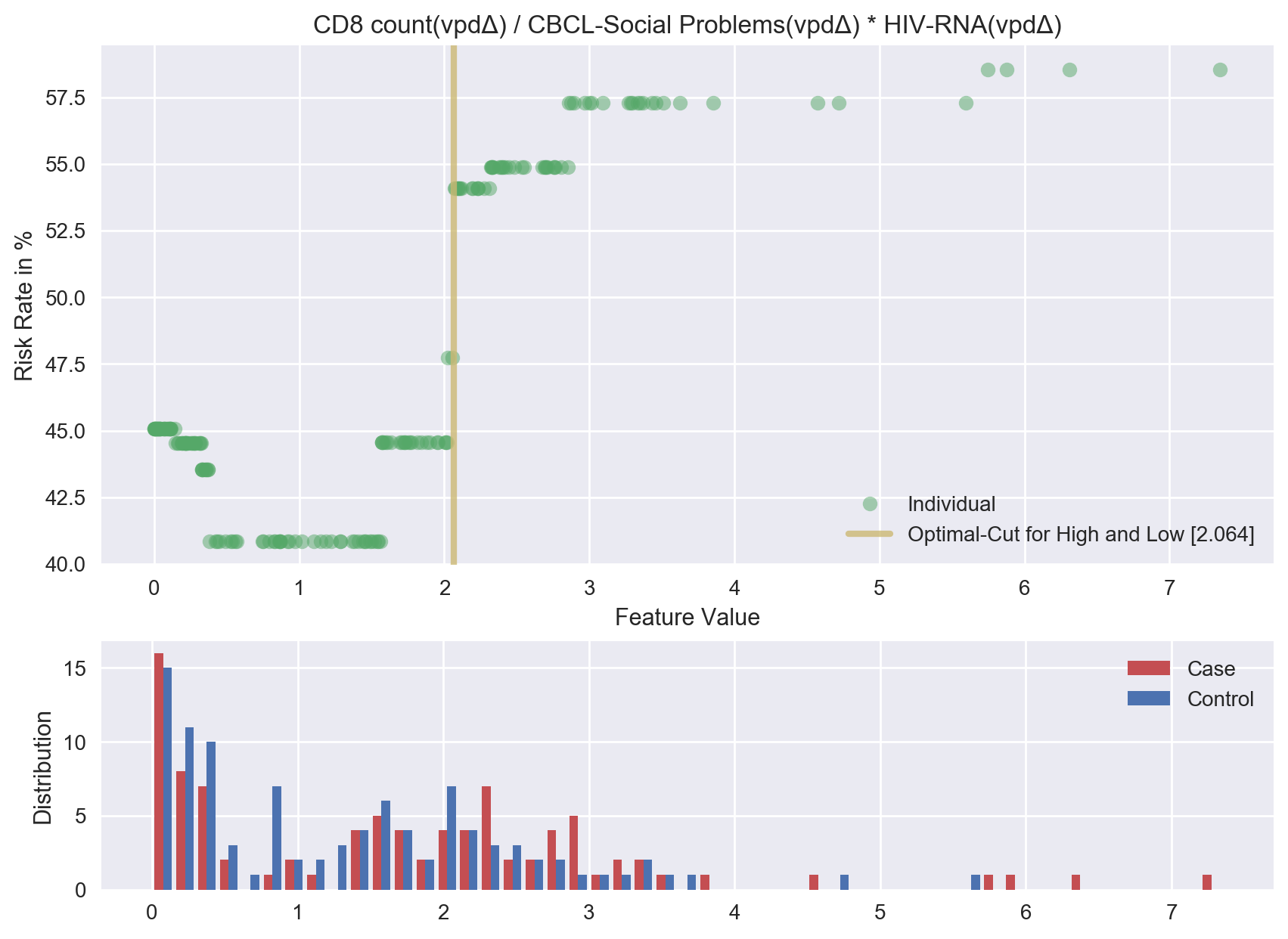

**Feature 24.Cluster of CD4 Percentage, CBCL Somatic Problems Score, and Mean Corpuscular Hemoglobin**

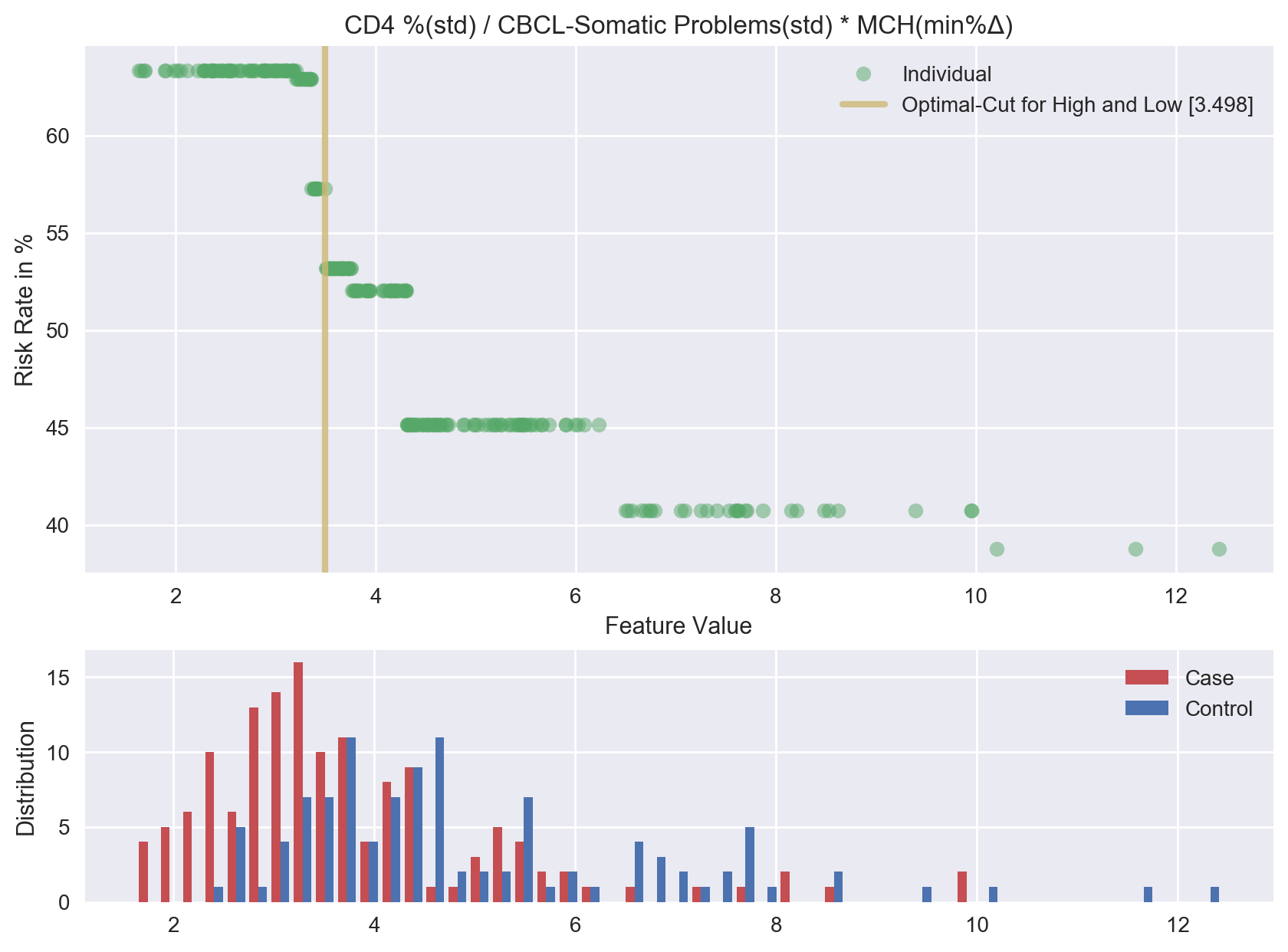

**Feature 25. Total CBCL Score**

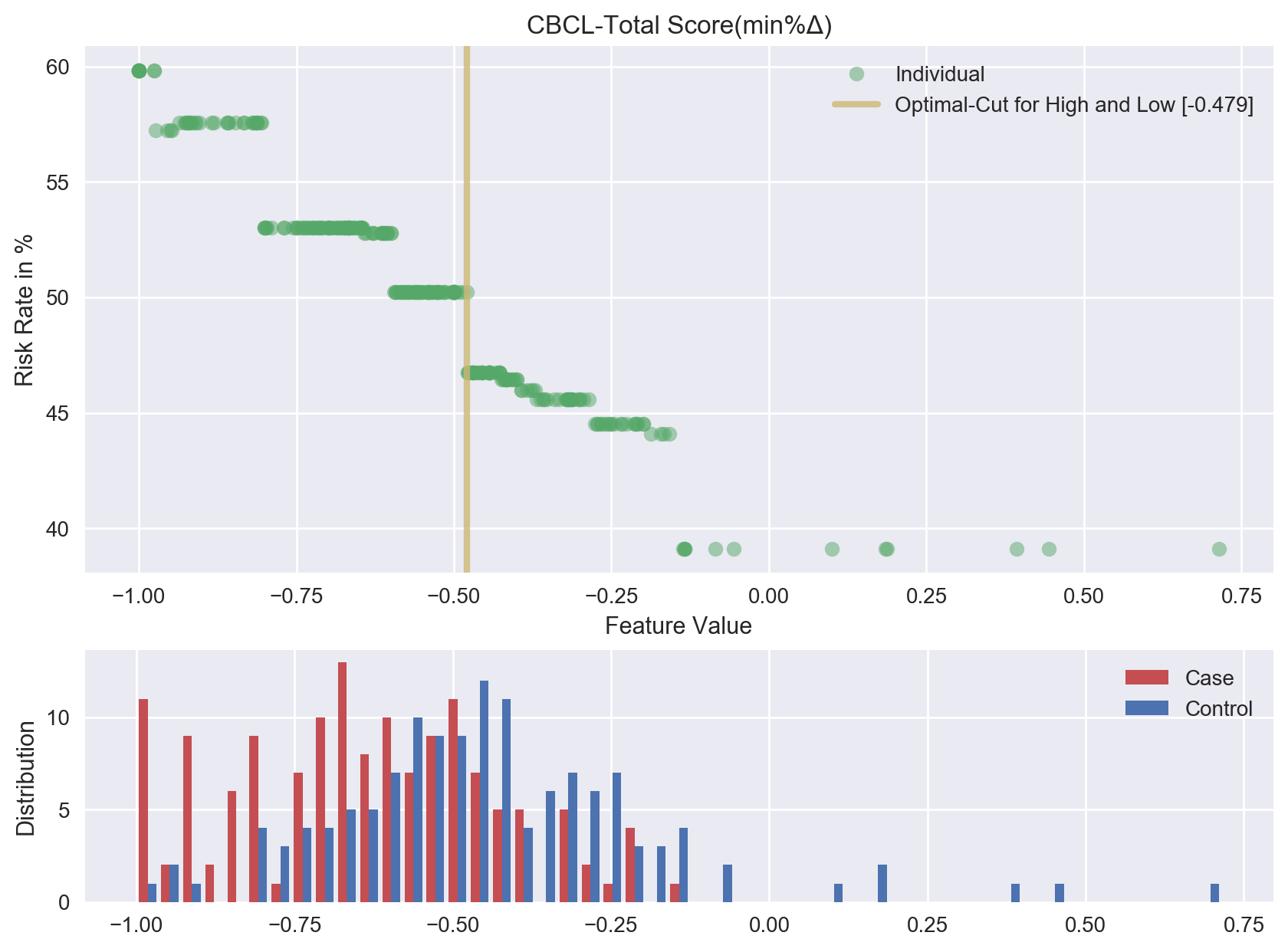

**Figure S2. Frequency of temporal change, dispersion and measures of central tendency in the predictive algorithm.**

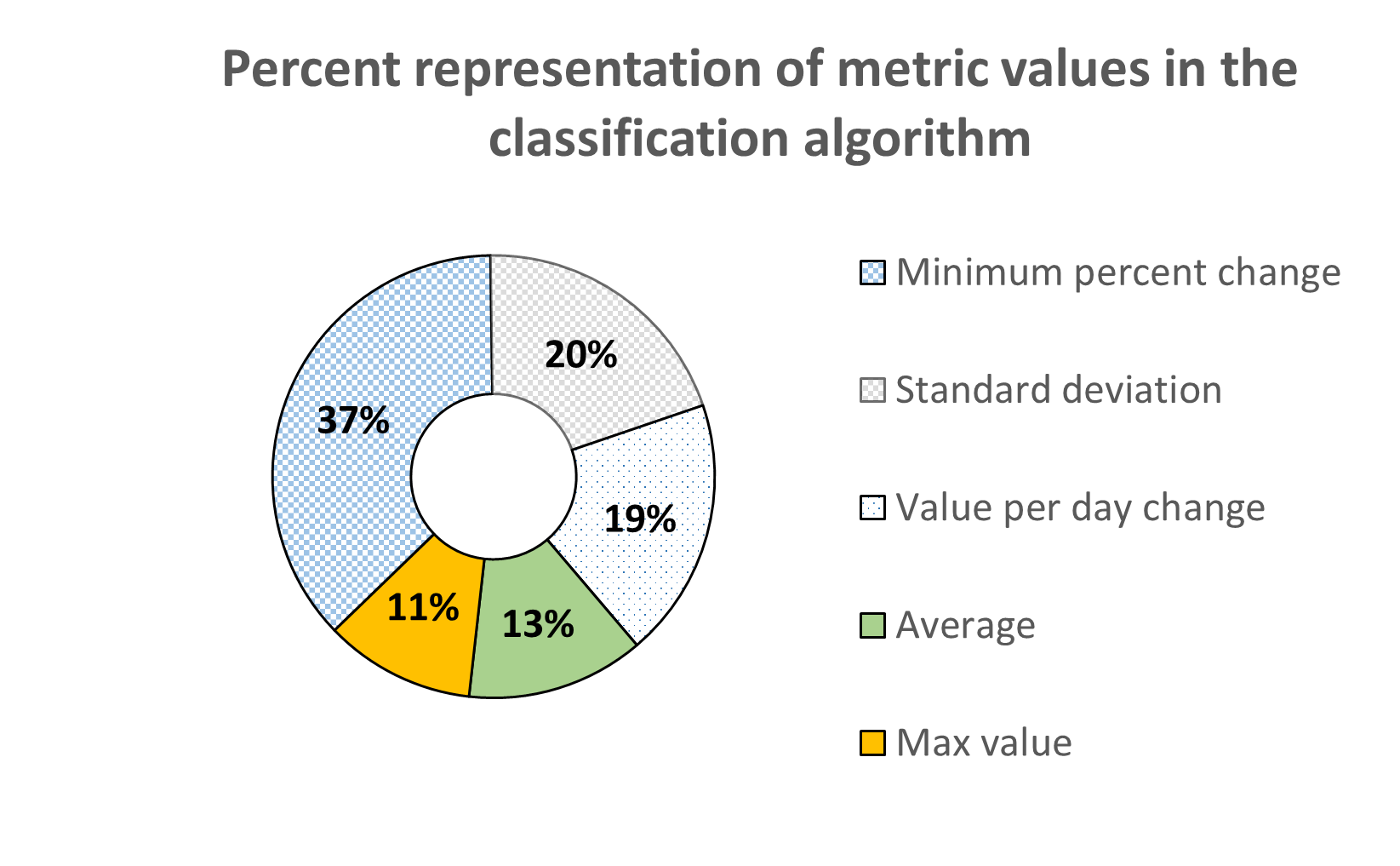

**Fig. S2.** Representation of each variable type in the final predictive model as defined by the machine learning analysis. Lighter colors (blue, gray, white) represent nontraditional metrics of change and dispersion; darker colors (gold, green) represent more common measures of central tendency. The nontraditional features identified by the machine learning analysis accounted for nearly 75% of the metrics selected by the machine learning analysis.

**Figure S3. Box plots for the top 10 predictive features.**

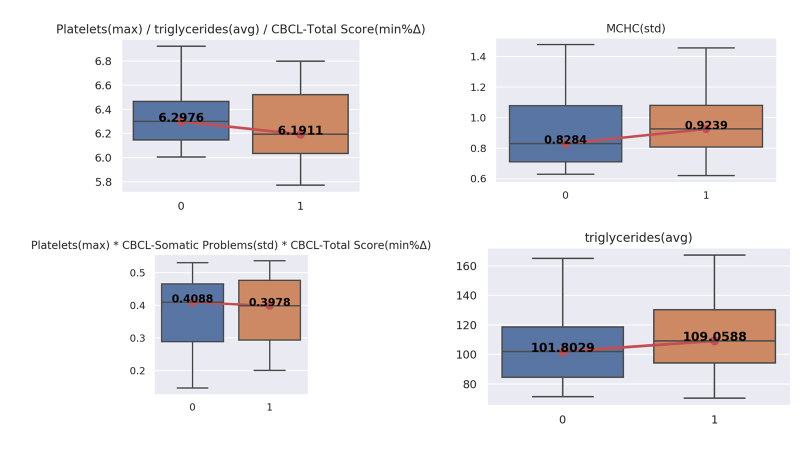

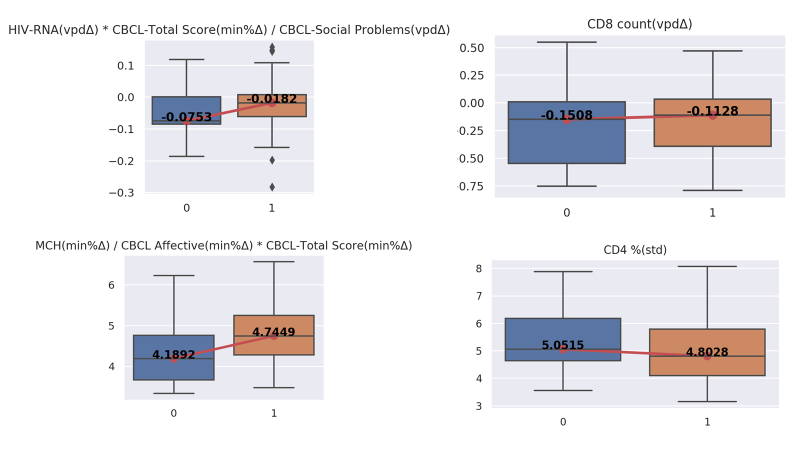

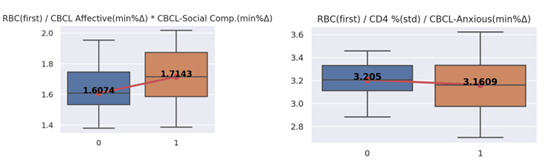

**Table S1**. **Frequency of each variable in the machine learning algorithm.**

| Variable | Frequency of each variable among the top features |
| --- | --- |
| CBCL-Total Score | 7 |
| Triglycerides md/dL | 5 |
| CBCL-Affective Scale | 5 |
| CBCL-Somatic Problems Scale | 4 |
| CBCL-Social Problems Scale | 4 |
| Plasma HIV-RNA viral load | 4 |
| CD4 T cell count percent | 4 |
| Mean corpuscular hemoglobin concentration g/dL | 3 |
| CBCL-Anxious Scale | 3 |
| Platelets 10^3/ul | 3 |
| CD8 T cell count cell/mm^3^ | 2 |
| Mean corpuscular hemoglobin concentration pg/dL | 2 |
| CBCL-Internalizing Problems Scale | 2 |
| Red blood cell count | 2 |
| CBCL-Social Competence Scale | 2 |
| Lymphocytes 10^3/ul | 1 |
| CBCL-Activities Competence Scale | 1 |

**Table S2. Logistic regression classification of neurocognitive trajectories.**

| **Predictor Variable** | **Odds Ratio** | **P>z** | **95% Confidence Interval** | **Interval** |
| --- | --- | --- | --- | --- |
| CD4/CD8 Ratio | 3.41 | 0.586 | 0.04 | 286.17 |
| HIV Viral Load | 0.99 | 0.462 | 0.99 | 1.00 |
| CBCL Total Score | 1.05 | 0.121 | 0.98 | 1.12 |
| Family Income | 3.44 | 0.393 | 0.20 | 59.10 |
| CD4/CD8 Ratio X HIV Viral Load | 0.99 | 0.535 | 0.99 | 1.00 |
| CD4/CD8 Ratio X CBCL Total | 0.88 | 0.074 | 0.78 | 1.01 |
| CD4/CD8 Ratio X Family Income | 0.37 | 0.699 | 0.00 | 55.42 |
| HIV Viral Load X CBCL Total Score | **1.00** | **0.028** | **1** | **1** |
| HIV Viral Load X Family Income | **0.99** | **0.018** | **0.99** | **0.99** |
| CBCL Total Score X Family Income | 1.00 | 0.678 | 0.97 | 1.03 |
| Constant | 0.99 | 0.998 | 0.09 | 10.83 |

Logistic regression Prediction of BVMI trajectory classification (higher/lower). HIV viral load x CBCL Total Score and HIV viral load x Family Income were significant predictors. Total accuracy of the model was 45% (Log likelihood = -175.20, Pseudo R2=0.1129; Chi^2^(10) = 44.60).
